# LC3-associated endocytosis facilitates extracellular Tau aggregate internalization and degradation in microglia

**DOI:** 10.1101/2024.12.06.627303

**Authors:** Hariharakrishnan Chidambaram, Subashchandrabose Chinnathambi

## Abstract

**Background:** Microglia, which are resident phagocytic cells in the brain, are involved in the active clearance of microbes, misfolded proteins, and cell debris, among others. In Alzheimer’s disease, microglia play a pivotal role in clearing extracellular amyloid-β (Aβ) plaques and intracellular Tau aggregates from the brain. Microglial cells have several mechanisms for Tau internalization, including macropinocytosis, heparan sulfate proteoglycans (HSPGs), dynamin-dependent endocytosis, and receptor-mediated endocytosis. Internalized Tau seeds either undergo proteasomal or lysosomal degradation or are exocytosed into the extracellular space via exosomes. Microtubule-associated protein 1 light chain 3 (LC3)-associated endocytosis (LANDO) is a recently discovered microglial mechanism for an effective clearance of Aβ aggregates and alleviates neurodegeneration in murine Alzheimer’s disease. Several microglial receptors are reported to be involved in misfolded Aβ and Tau aggregate internalization such as triggering receptor expressed on myeloid cell 2 (TREM-2), P2Y purinoceptor 12 (P2Y12R), C- X3-C motif chemokine receptor 1 (CX3CR1), etc.

**Methods:** In this study, we report the LANDO of Tau monomers and aggregates by murine microglial cells, which are further degraded by lysosomal fusion. LANDO of extracellular Tau is demonstrated by several biochemical and cell-biology studies including western blotting, fluorescence and confocal imaging of microglial cells.

**Results:** We analyzed microglial activation by measuring the upregulation of Ionized Calcium-binding Adapter Molecule 1 (Iba-1) expression. Later, we demonstrated the LANDO of human full-length Tau species, where LC3 colocalizes with internalized Tau, followed by lysosomal degradation by microglia. The accumulation of internalized Tau in the perinuclear region of the cell in the presence of chloroquine supports phagolysosomal fusion and degradation.

**Conclusion:** Hence, we concluded that microglia internalize extracellular Tau species via LANDO, which further undergoes degradation via the lysosomal pathway.

## Introduction

Microglia are the resident phagocytic cells that are involved in the active clearance of microbes, apoptotic cells, and misfolded protein aggregates such as Tau, amyloid-β (Aβ), and cell debris in the brain environment.(^1–3^) Microglia are involved in constant surveillance in their ramified form (inactive), which activates (amoeboid), proliferates, and migrates toward the site of injury.(^2, 4^) Microglial cells are activated by a wide variety of signal molecules such as nucleotides, peptides, and chemokines.(^5^) The signal is received through surface receptors such as G-protein coupled receptors (GPCRs), ion channels, and transmembrane receptors to attain either a pro- (M1) or anti-inflammatory (M2) microglial state.(^6–8^)

Tau is a microtubule-associated protein expressed in neurons that plays a pivotal role in microtubule stabilization and cargo trafficking.(^9–11^) Under pathological conditions, Tau undergoes several post-translational modifications, such as phosphorylation, acetylation, sumoylation, and truncation, that detach it from the microtubules to form oligomeric and aggregated structures.(^12–14^) In the brains of patients with Alzheimer’s disease (AD), accumulated Tau from the intracellular space of neuronal cells is released into the extracellular environment by several mechanisms, which implant a seeding effect and spread to neighboring healthy neuronal cells.(^15^) Tau seeds are spread either from leaky substances of dying neurons or secreted into the extracellular space via vesicles called exosomes to alleviate Tau-induced toxicity in cells.(^16–19^) Tau propagation to adjacent neurons and glia occurs through several mechanisms, such as clathrin-mediated endocytosis, receptor- or heparan sulfate proteoglycan (HSPG)- mediated endocytosis, and nanotubes, among others.(^3, 20^) In microglia, extracellular Tau is internalized via a HSPG-mediated mechanism, which is either accumulated in the cytosol as seeds, exocytosed through vesicles, or degraded by lysosomal fusion.(^21, 22^) Monomeric Tau has been reported to be internalized by micropinocytosis, whereas aggregated Tau is internalized via dynamin-dependent pathways that are enveloped in endosomes for degradation.(^23^) Tau is also reported to be internalized by microglial membrane receptors such as P2Y purinoceptor 12 (P2Y12R) and C-X3-C motif chemokine receptor 1 (CX3CR1).(^24–32^)

Phagocytosis is the engulfment of extracellular materials by extended actin-rich structures called phagocytic cups, which result in the internalization of the material into a single membrane vesicle (phagosome), ultimately leading to degradation by fusion with lysosomes.(^33–35^) Specific immune cells, dendritic cells, monocytes, and macrophages phagocytose invading microbes, extracellular debris, and protein aggregates. Under AD conditions, microglial cells undergo Tau endocytosis, which is available in the extracellular brain environment as monomers, oligomers, and neurofibrillary filaments.(^24, 36–38^) Autophagy is the clearance of intracellular microbes, organelles, cytosolic substances, and protein aggregates by sequestering in a double membrane-bound structure called autophagosome.(^39, 40^) Microtubule-associated protein 1 light chain 3 (LC3), a mammalian homolog of autophagy-related protein (ATG) 8 autophagy protein, is a specific and widely used marker for determining autophagy influx in cells.(^41, 42^) Autophagosome formation is directly related to LC3-II levels in cells.(^43^) During autophagy, the autophagy protein ATG4 cleaves the C-terminus of LC3 to form LC3-I. LC3-I is then converted to LC3-II by conjugation with lipid phosphatidylethanolamine, which binds to the outer and inner membranes of the autophagosome.(^44^) The LC3-II associated autophagosomes containing sequestered cytosolic materials are ultimately associated with lysosomes for degradation by forming autolysosomes.(^45, 46^) LC3-associated phagocytosis (LAP) is a non-canonical autophagy mechanism that phagocytoses extracellular cargo, such as apoptotic cells, Toll-like receptor (TLR) ligand-coated latex beads, zymosans, and microbes.(^47–50^) In LAP, LC3-II is recruited to the phagosomal membrane upon phagocytosis and promotes its trafficking to fuse with lysosomes for active degradation.(^51^) LAP enhances phagolysosomal fusion, which determines its anti-microbial activity.(^51^) Similarly, LC3-associated endocytosis (LANDO) of Aβ by microglial cells in murine AD has been recently reported to protect from Aβ toxicity, neuronal loss, and memory impairment.(^52^) In tauopathies, such as familial AD, accumulation of autophagic and lysosomal markers, LC3, and lysosome-associated membrane protein (LAMP) 1, was observed, indicating defects in the autophagosomal and lysosomal pathways, respectively.(^53^) The levels of LC3-positive vesicles increased in the frontal cortex of patients with AD, which colocalizes with Tau, indicating the accumulation of autophagic vesicles.(^53^) In this study, we demonstrated the involvement of LANDO in Tau internalization using confocal fluorescence microscopy in murine microglial cells. We demonstrated microglial activation and LANDO, followed by lysosomal degradation. Chloroquine is an antimalarial drug that impairs lysosomal degradation by autophagosome-lysosomal fusion.(^54, 55^) We observed LC3-associated endosomal accumulation along with internalized Tau in chloroquine-treated groups to demonstrate its lysosomal degradation. This study clearly demonstrates LANDO and degradation of extracellular Tau, suggesting a molecular mechanism for its clearance in AD.

## Methods

### Preparation of Tau

Tau protein was prepared according to the protocol described by Chidambaram and Chinnathambi(^56^). The methodology is briefly described below. Human full-length Tau (hTau40) was recombinantly expressed in the BL21 strain of *Escherichia coli* cells. The cells were grown at 37 with constant shaking until the optical density (OD) reaches 0.5–0.6. Tau protein expression is induced with 0.5 mmol/L isopropyl β-d-1-thiogalactopyranoside (IPTG) and further incubated for 4 h. The cells were then pelleted and resuspended in a cell lysis buffer. 2- (N-morpholino) ethanesulfonic acid (MES) buffer pH 6.8 was used for cell lysis that includes 20 mmol/L MES, 1 mmol ethylene glycol-bis (baminoethylether)-N,N,N′,N′-tetraacetic acid (EGTA), 0.2 mmol/L MgCl_2_, 5 mmol/L dithiothreitol (DTT), 1 mmol/L phenylmethylsulfonylfluoride (PMSF), and protease inhibitor complex (PIC) (0.01%). The cells were homogenized at 15,000 psi in a cell disruption system and added with 0.5 mol/L NaCl and 5 mmol/L DTT were added to boil the lysate (90 for 15 min). This step allows protein precipitation of lysate except for a few highly soluble proteins, such as Tau, that are separated (from supernatant) by centrifuging the lysate at 40,000 rpm for 45 min at 4 . The excess salt from the lysate is then removed by dialyzing the supernatant with lysis buffer (with 5 mmol/L NaCl) overnight at 4 and centrifuged again at 40,000 rpm for 45 min for further precipitation. Tau protein was separated from the supernatant by cation-exchange chromatography with gradient elution using 1 mol/L NaCl. The protein was further purified by size exclusion chromatography in PBS buffer containing 2 mmol/L DTT and concentrated using 5 kDa cutoff centricons. The monomeric Tau protein concentration was estimated using the bicinchoninic acid (BCA) assay, aliquoted in multiple tubes, snap-frozen, and stored at 80 for further studies.

Tau aggregates are prepared by incubating Tau protein (100 µmol/L) with heparin 17,500 Da (25 µmol/L) at a ratio of 4:1 at 37 for 72 h. The aggregates were prepared in 20 mmol/L N,N- bis(2-hydroxyethyl)-2-aminoethanesulfonic acid (BES) buffer (pH 7.4) supplemented with 1 mmol/L DTT, 0.01% sodium azide, 25 mmol/L NaCl, and 10% PIC. The quality of the Tau monomers and aggregates was determined using sodium dodecyl sulfate (SDS) gel electrophoresis, Thioflavin-S (ThS) fluorescence assay, and transmission electron microscope (TEM) analysis (^57–62^).

### Labeling of Tau monomer and aggregate

hTau40 (100 µmol/L) is added with tris(2-carboxyethyl)phosphine (TCEP) (2 µmol/L) followed by the gradual addition of Alexa Fluor^647^ C_2_ maleimide in the ratio of 2:1 (C_2_ maleimide: Tau) and incubated overnight at 4 shaker. Buffer exchange was performed by adding excess phosphate buffered saline (PBS, pH 7.4) using 5-kDa centrifugal filters, and the final concentration was estimated using the BCA assay. The quality of the Alexa Fluor^647^ labeled Tau monomer and aggregates was determined using SDS-gel electrophoresis, ThS fluorescence assay, and TEM analysis.

### Culturing of N9 microglial cells

N9 microglial cells (RRID:CVCL_0452) were cultured in Roswell Park Memorial Institute (RPMI) medium supplemented with 10% fetal bovine serum (FBS) and incubated at 37 with 5% CO_2_. The cells were harvested at 80% confluence and seeded for immunofluorescence and Western blot analyses.

### Western blot

The expressions of different proteins upon 1 µmol/L Tau treatment are studied using Western blot analysis. N9 cells were counted, and three lakh cells per well were seeded in multi-well dishes. Cells were incubated for 24 h at 37 with 5% CO_2_ supply. The medium was drained, and the cells were incubated with reduced serum medium containing 0.5% FBS. Then, 1 µmol/L Tau (monomer and aggregate) treatment is then provided and incubated for 24 h along with control group (no Tau treatment). Cells were harvested by trypsinization with 0.05% trypsin-EDTA (1X), washed with PBS (pH 7.4), and pelleted by centrifugation at 800 rpm for 5 min. The cells were lysed with 100 µL RIPA buffer containing 1.0% Triton X-100 and 10% PIC and centrifuged to pellet down the nucleus and membrane particles at 14,500 rpm for 20 min. The protein concentration in the supernatant containing soluble proteins was estimated using Bradford assay. An equal protein load (75 µg per sample) was used for Western blot analysis. The samples were subjected to SDS gel electrophoresis and transferred to a nitrocellulose membrane activated with methanol for 5 min. The transfers were performed using an Amersham semi-dry transfer unit at 200 mA. The membrane containing the transferred proteins was blocked with 5% skim milk in PBS containing 0.1% Tween-20. The membrane is then incubated with primary antibody followed by secondary antibodies with the mentioned dilutions: Iba-1 rabbit polyclonal antibody, 1:1000; LC3 polyclonal antibody, 1:500; and β-actin loading control monoclonal antibody, 1:2500. The blot is developed at different exposure times with SuperSignal™ West Atto Ultimate Sensitivity Substrate. The band intensity was quantified using Image Lab software version 6.0.1 (Bio-Rad, Gurugram, India) and normalized using β-actin as a housekeeping gene. The experiment was analyzed using three independent repeats (*n* = 3), and the fold change in the expression of Iba-1 and LC3 was studied (^25^).

### Immunofluorescence assay

Tau internalization and the localization of Iba-1, LC3, and lysosome-associated membrane protein-2A (LAMP2A) with F-actin were studied via immunofluorescence microscopy. N9 cells were counted and 10,000 cells per well were seeded onto coverslips placed in multi-well dishes. Cells are incubated for 24 h at 37 with a 5% CO_2_ supply. The medium was drained, and the cells that adhered to the coverslips were provided with reduced serum medium containing 0.5% FBS. Then, 1 µmol/L Alexa Fluor^647^-tagged Tau treatment (monomer and aggregates) is provided and incubated for 24 h along with the control group (no Tau treatment). In inhibitor-based study, chloroquine (30 µmol/L) is provided as a 30-min pre-treatment. Alexa Fluor^647^-tagged Tau treatment (1 µmol/L, monomer and aggregates) is then provided and incubated for 3 h along with Tau control (only Tau treatment) and blank control (no Tau treatment). The cells were fixed with 4% paraformaldehyde for 15–20 min at room temperature and coverslips containing the cells were washed with PBS. The cells were permeabilized with PBS containing 0.2% Triton X (PBS-t) for 20 min and blocked with PBS-t buffer containing 5% horse serum. Finally, cells were incubated with the respective primary (4 overnight) and secondary antibodies (1 h at room temperature) at the following dilutions: Iba-1, 1:200; LC3, 1:400; LAMP2A, 1:500; and Alexa Fluor^488^ tagged phalloidin (F-actin), 1:50, followed by nuclear staining with 4’,6-diamidino-2-phenylindole (DAPI, 300 nmol/L) for 10 min. The coverslips were mounted with Prolong Diamond Antifade mountant and dried for 24 h at room temperature. Images of N9 microglial cells were captured using a Zeiss Axio observer Z1/7 Apotome 2.0 wide-field fluorescence microscope (Bangalore, India) and a Leica Stellaris 5 confocal microscope (Mumbai, India). Plan Apochromat 63x/1.40 Oil DIC M27 objective lens and an Axiocam 503 camera were used for visualization and image capture, respectively, using a fluorescence microscope. Plan Apochromat 100x/1.40 objective and hybrid detector S (HyD S) silicon detectors were used for visualization and image capturing, respectively, in a confocal microscope. The post-capture processing of two-dimensional and Z-stack images, followed by intensity quantification, was performed using Zen 2.3 (Zeiss, Bangalore, India) and Leica Application Suite X 4.5 (Leica Biosystems, Mumbai, India) software. Pearson’s co-efficient of colocalization between Iba-1, LAMP2A, and F-actin was analyzed by using ImageJ software (Bethesda, Maryland, USA) with Coloc2 plugin at a different field of interest from all treatment groups as compared to the control group (*n* ≥15).

### Statistical analysis

All experiments were performed in three biological replicates, and triplicate measurements are taken for each experiment. Statistical significance was determined using one-way analysis of variance (ANOVA) using GraphPad Prism 8.0.1 software (Boston, Massachusetts USA, www.graphpad.com). For multiple groups, significance was calculated using Tukey–Kramer’s post hoc analysis at a 5% level of significance. The *P* values are then calculated by comparing with control group, and the level of significance is mentioned within the graph (*** *P* <0.001, ** *P* <0.01, * *P* <0.05, ^ns^ *P* ≥0.05). The results are considered significant if the mean difference between treatment groups is greater than calculated Tukey’s criterion (*XX’>T)*.

## Results

### Extracellular Tau activates microglia and promotes Tau internalization

Human Tau is a 441 amino acid protein that self-aggregates to form higher-order species in the AD brain.(^56^) Human recombinant full-length Tau forms AD-like paired helical filaments *in vitro* in the presence of several anionic cofactors such as heparin and ribonucleic acid (RNA), etc.(^56, 63–68^) In this study, we purified human recombinant Tau and induced *in vitro* to form aggregates with heparin.(^56^) Recombinant full-length Tau monomers and aggregates were characterized using biophysical and biochemical studies and labeled with Alexa Fluor^647^ C2- maleimide that conjugates to the thiol group of target proteins (Supplementary figure. 1). Tau activates microglia by directly binding to several receptors such as polyglutamine-binding protein 1 (PQBP1) and purinergic receptors such as the P2Y12 receptor.(^25, 69^) Ionized Calcium-binding Adapter Molecule 1 (Iba-1) is a 17 kDa molecule that binds to filamentous actin (F-actin) structures at membrane ruffles and phagocytic cups upon activation of macrophages.(^70, 71^) We studied the levels and distribution of the Iba-1 protein upon extracellular Tau treatment for microglial activation and migration [Figure 1]. N9 microglial cells stained with Iba-1 and phalloidin (F-actin) demonstrated increased Iba-1 intensity levels upon monomer and aggregate treatment for 24 h compared to the control group [Figures 1A and 1C]. Iba-1 colocalizes with F-actin at the membrane ruffles and at the leading edge (lamellum) upon Tau monomer treatment, followed by aggregates, where no significant colocalization was observed in control group denoting microglial activation and endocytosis [Figures 1A and 1B]. Membrane ruffling and Iba- 1 colocalization in the lamellum were clearly visualized using an orthogonal representation [Figure 1B]. The degree of colocalization was quantified using Pearson’s coefficient [Figure 1D]. Iba-1 expression increased upon extracellular Tau treatment for 24 h, as determined by Western blot [Figure 1E] and the corresponding quantification [Figure 1F]. These data suggest that Tau-mediated microglial activation and endocytosis occurs via membrane ruffling [Figure 1G].

**Figure 1:**
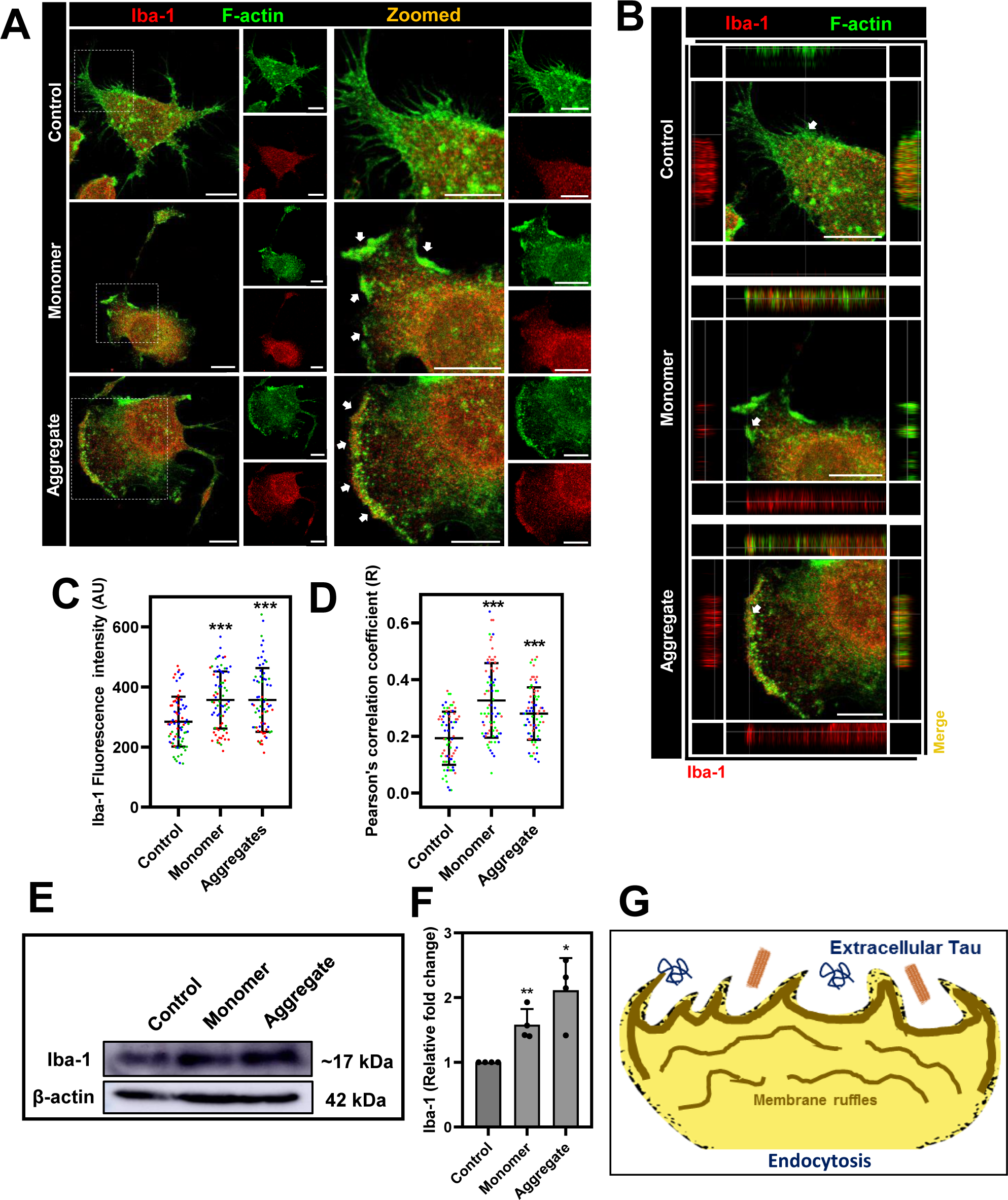
Extracellular Tau activates microglia for internalization. (A) Confocal microscope images of N9 microglial cells stained with Iba-1and phalloidin (stains filamentous actin [F- actin]) showing colocalization in monomer (1 µmol/L) and aggregate (1 µmol/L)-treated groups. F-actin staining visualizes membrane ruffling (indicated by white arrows in Tau-treated groups) and active colocalization with Iba-1 at the lamellipodial surface of Tau-treated microglial cells. Scale bar represents 10 µm. (B) Orthogonal representation of microglial that reveals membrane ruffling and Iba-1 colocalization at the lamellum of Tau-treated cells. The left and bottom axis reveals Iba-1, whereas the right and top axis represent colocalization between F-actin and Iba-1 at membrane ruffles for better visualization. (C) Intensity mean values of Iba-1 quantified and plotted from 25 different cells from each biological repeat demonstrating increased levels of Iba- 1 (each color code represents a biological repeat) (*n* = 75). The data is statistically significant with a *P* value <0.05 calculated using one-way ANOVA, and the statistical significance of the multiple groups has been calculated using Tukey–Kramer’s post hoc analysis for multiple comparisons at 5% level of significance. (D) Pearson’s co-efficient of colocalization between Iba-1 and F-actin (each color code represents a biological repeat) (*n* = 75). The data is statistically significant with a *P* value <0.05 calculated using one-way ANOVA and the statistical significance of the multiple groups has been calculated through Tukey–Kramer’s post hoc analysis for multiple comparisons at 5% level of significance. (E and F) The expression levels of Iba-1 upon extracellular Tau treatment for 24 h were determined by Western blot analysis and the corresponding quantification (*n* = 4). The data is statistically significant calculated by ratio-paired Student’s *t*-test and compared with the control group. (G) Schematic representation of microglial activation and endocytosis by actin-associated membrane ruffling. *** *P* <0.001, ** *P* <0.01, * *P* <0.05, ^ns^ *P* ≥0.05. ANOVA: Analysis of variance; F-actin: Filamentous actin; Iba-1: Ionized Calcium-binding Adapter Molecule 1..

### Microtubule-associated protein 1 light chain 3-associated endocytosis of extracellular Tau by microglia

Upon sensing extracellular foreign particles, microglia form cup-like actin-rich phagocytic structures for active internalization.(^36, 37, 72, 73^) Several microglial surface receptors such as P2Y12R and CX3CR1 are involved in misfolded Tau protein interactions and internalization.(24–26, 74) Tau internalization by clathrin-mediated endocytic pathways is a well-studied mechanism that promotes vesicular trafficking of Tau through early and late endosomes for lysosomal degradation.(^37, 75^) To study LANDO of extracellular Tau species, microglial cells were treated with extracellular monomeric and aggregated Tau species and analyzed for Tau endocytosis by staining with LC3 and F-actin [Figure 2 and Supplementary figures 2 and 3, cells focused on endocytic structures]. Tau monomers and aggregates were provided extracellularly, and the cells were studied at different time intervals (3, 6, 12, and 24 h) for Tau internalization. Actin-rich endocytic structures were observed in the monomers [Figure 2A] and aggregate-treated groups [Figure 2B] irrespective of the time interval. Endocytic structures containing extracellular Tau species formed at the leading edge of microglia stained with F-actin and colocalized with LC3 in the Tau-treated groups. We resolved these structures using a Stellaris 5 confocal microscope to observe the association between LC3-II and endosomal membranes [Figure 3A and Supplementary figure 4]. The endosomal structures (marked by arrows) observed in the magnified images (right) clearly show an association of LC3. The corresponding Z-stack images of Tau-containing endosomes from the cortical to the internal plane show co-localization between actin filaments, LC3, and internalized Tau [Figure 3B]. Pearson’s coefficient showed co-localization between Tau and LC3 upon endocytosis in Tau-treated groups, supporting the association of Tau in LC3-associated endosomes [Figure 3C]. The cortical LC3 intensity levels quantified from the immunofluorescence study increased in the Tau-treated groups compared to those in the control group [Figure 3D]. To support this, we studied the LC3 levels, as detected by Western blot analysis, and found no significant change in expression upon extracellular Tau treatment compared to the control group [Figures 3E and 3F]. This suggests a cortical association of LC3 in the Tau-treated groups [Figure 3D], as there was no change in total expression. The LC3-II fold-change (lipid phosphatidylethanolamine-conjugated LC3 (represents the lower band in Figure 3E) significantly increased in the monomer- and aggregate-treated groups, as shown in Figure 4G denoting LC3-II conjugation to the endosomal membranes. These results suggest the involvement of LC3-II in Tau endocytosis by microglial cells, as is evident from the colocalization of LC3, F-actin, and Tau upon microglial activation [Figure 3H].

**Figure 2:**
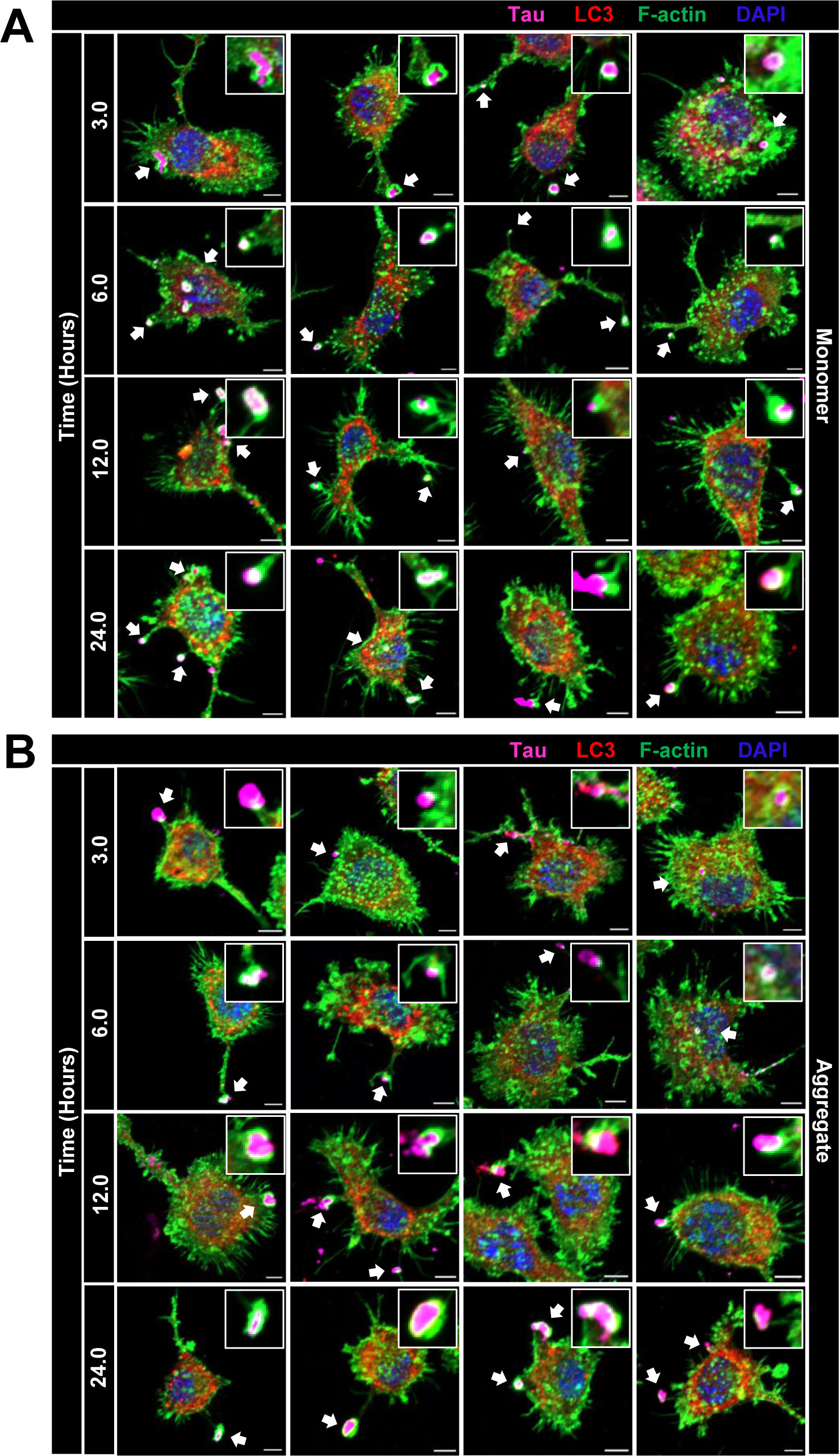
Microglial endocytic structures for Tau internalization. (A) Microglial endocytic structures observed for Tau monomer internalization at all-time intervals (The small figures within individual figure shows the zoomed endocytic structures containing Tau colocalizing with LC3 and F-actin). (B) Microglial endocytic structures observed for Tau aggregate internalization at all-time intervals (The small figures within individual figure shows the zoomed endocytic structures containing Tau colocalizing with LC3 and F-actin). Scale bar represents 5 µm. F-actin: Filamentous actin; LC3: Microtubule-associated protein 1 light chain 3.

**Figure 3:**
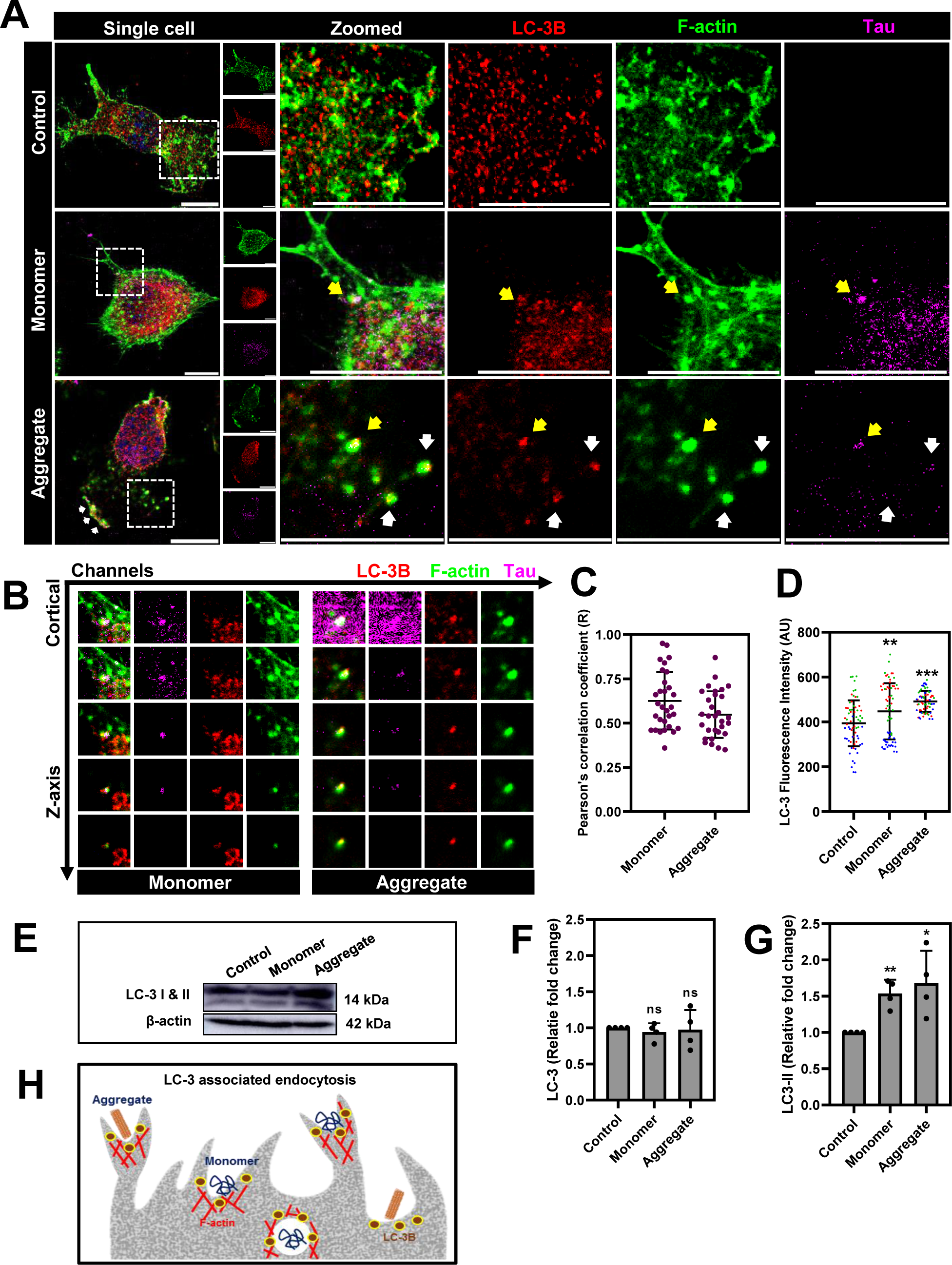
LC3-associated endocytosis of extracellular Tau. (A) Confocal microscope images of N9 microglial cells treated with Alexa Fluor^647^-tagged monomeric and aggregated Tau for 3 h stained with LC3 and phalloidin (F-actin). The two-dimensional representation of microglial cells internalizing extracellular Tau that colocalizes with LC3 and F-actin acquired by Leica Stellaris 5 confocal microscope (arrows indicate the endocytic structure in Tau-treated groups). The scale bar represents 10 µm. (B) Z-stack representation of Tau-containing endosomes in microglia (obtained from endosomes visualized in Figure 4A marked with yellow arrows). Z-stack images are captured at an interval of 0.24 µm from the cortical toward internal plane (z-axis), which reveals a three-point colocalization between Tau, LC3, and F-actin. (C) Pearson’s co-efficient showing colocalization of Tau and LC3 in microglial endosomes (*n* = 30) (region of interest selected only from endocytic structures). (D) Quantification of cortical LC3 intensity from immunofluorescence microscopy (each color code represents a biological repeat) (*n* = 60). The data is statistically significant with a *P* value <0.05 calculated using one-way ANOVA, and the statistical significance of the multiple groups has been calculated by Tukey–Kramer’s post hoc analysis for multiple comparisons at 5% level of significance. (E) The expression levels of LC3-I and LC3-II examined by Western blot analysis reveal the active role of LC3-associated endocytosis and the corresponding quantification. (F) LC3 (I and II) relative fold change quantified from Western blot (*n* = 4. The data is statistically insignificant calculated by ratio-paired Student’s *t*-test and compared with the control group. (G) Relative fold change of LC3 II quantified from Western blot (lower band) (*n* = 4). The data is statistically significant with a *P* value <0.05 calculated using ratio-paired Student’s *t*-test and compared with the control group. (H) Summary of LC3 associated endocytosis of Tau species by microglial cells. *** *P* <0.001, ** *P* <0.01, * *P* <0.05, ^ns^ *P* ≥0.05. ANOVA: Analysis of variance; F-actin: Filamentous actin; LC3: Microtubule-associated protein 1 light chain 3.

**Figure 4:**
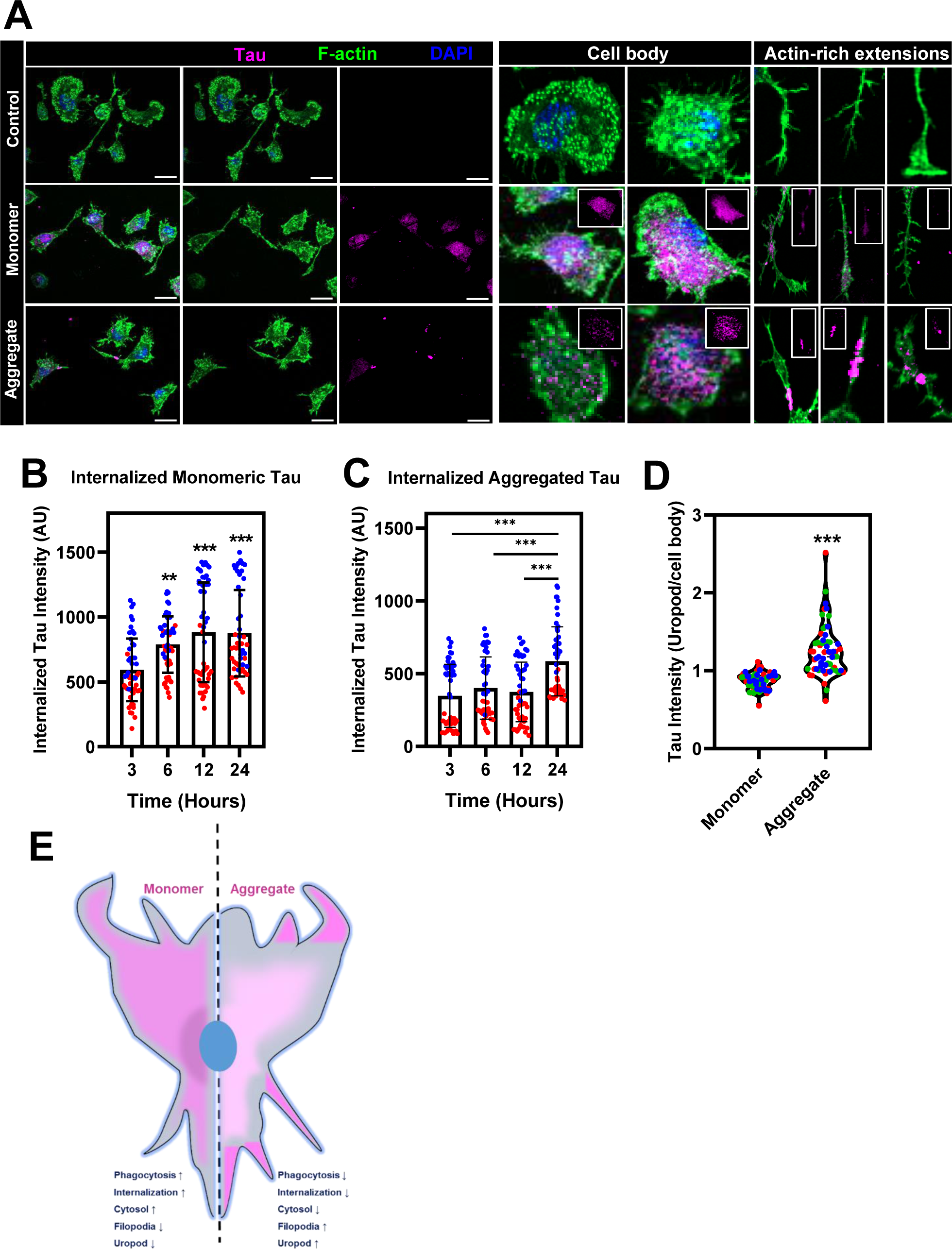
Internalized Tau monomer and aggregates are localized at different cytosolic locations. (A) Immunofluorescence analysis of N9 microglial cells treated with Alexa Fluor^647^- labeled monomeric and aggregated Tau for 24 h are stained with phalloidin (F-actin). The scale bar represents 20 µm. (B and C) Quantification of internalized Tau intensity for monomeric and aggregated Tau treatment at different time intervals (each color code represents a biological repeat) (*n* = 50). The data is statistically significant with *P* value <0.05 calculated using one-way ANOVA, and the statistical significance of the multiple groups has been calculated through Tukey–Kramer’s post hoc analysis for multiple comparisons at 5% level of significance. (D) Tau intensity ratio of filopodia to cell body quantified from monomer and aggregates treated microglial cell groups (each color code represents a biological repeat) (*n* = 60). The data is statistically significant with *P* value <0.05 calculated by unpaired Student’s *t*-test. (E) Diagrammatic representation of internalized Tau localization by microglial cells. *** *P* <0.001, ** *P* <0.01, * *P* <0.05, ^ns^ *P* ≥0.05. ANOVA: Analysis of variance; F-actin: Filamentous actin.

### Microglia localizes internalized Tau monomer and aggregate at different cytosolic locations

Furthermore, the distribution of monomeric and aggregated Tau along the cortical layer was visualized in different cells of the cell body and uropods, clearly showing the cytosolic distribution of monomeric Tau. Conversely, aggregates were mainly observed in actin-rich structures, such as filopodia and uropods [Figure 4A]. Next, we studied the internalized Tau intensity at different time intervals (3, 6, 12, and 24 h) using an immunofluorescence assay of monomer- and aggregate-treated cells. The mean intensity values of internalized Tau at different time intervals gradually increased with time, and the intensity values of monomeric Tau were comparatively higher than those of aggregated Tau. The intensity of the monomer was approximately 700 AU, whereas that of the aggregate reached approximately 500 AU after 24 h of Tau treatment [Figures 4B and 4C]. The ratio of Tau intensity in actin-rich structures to the cytosol was compared between the monomer- and aggregate-treated groups, calculated from the mean intensity values of internalized Tau at different cellular localizations [Figure 4D]. These data suggest time-dependent internalization of extracellular Tau species by microglial cells. Monomeric Tau is widely distributed throughout the cytosol, whereas aggregated Tau mainly accumulates in actin-rich structures, such as uropods and filopodia, which may either accumulate in the cells or further undergo degradation [Figure 4E].

### Lysosomal degradation of extracellular Tau species by microglia

The accumulated and misfolded proteins in cytosol undergo either ubiquitin-proteasomal or lysosomal degradation.(^76, 77^) We studied the lysosomal degradation of internalized Tau monomers and aggregates by immunofluorescence using a lysosomal marker, LAMP2A. Cytosolic monomeric Tau was highly colocalized with LAMP2A, followed by aggregates [Figure 5A]. The orthogonal representation of microglial cells showing the co-localization of internalized Tau with lysosomes for degradation [Figure 5B]. The level of Tau colocalization with LAMP2A was quantified by Pearson’s coefficient, showing comparatively higher values for monomers than for aggregates [Figure 5C]. These results suggest lysosomal degradation of the extracellular monomeric and aggregated Tau species, following the internalization by active endocytosis [Figure 5D].

**Figure 5:**
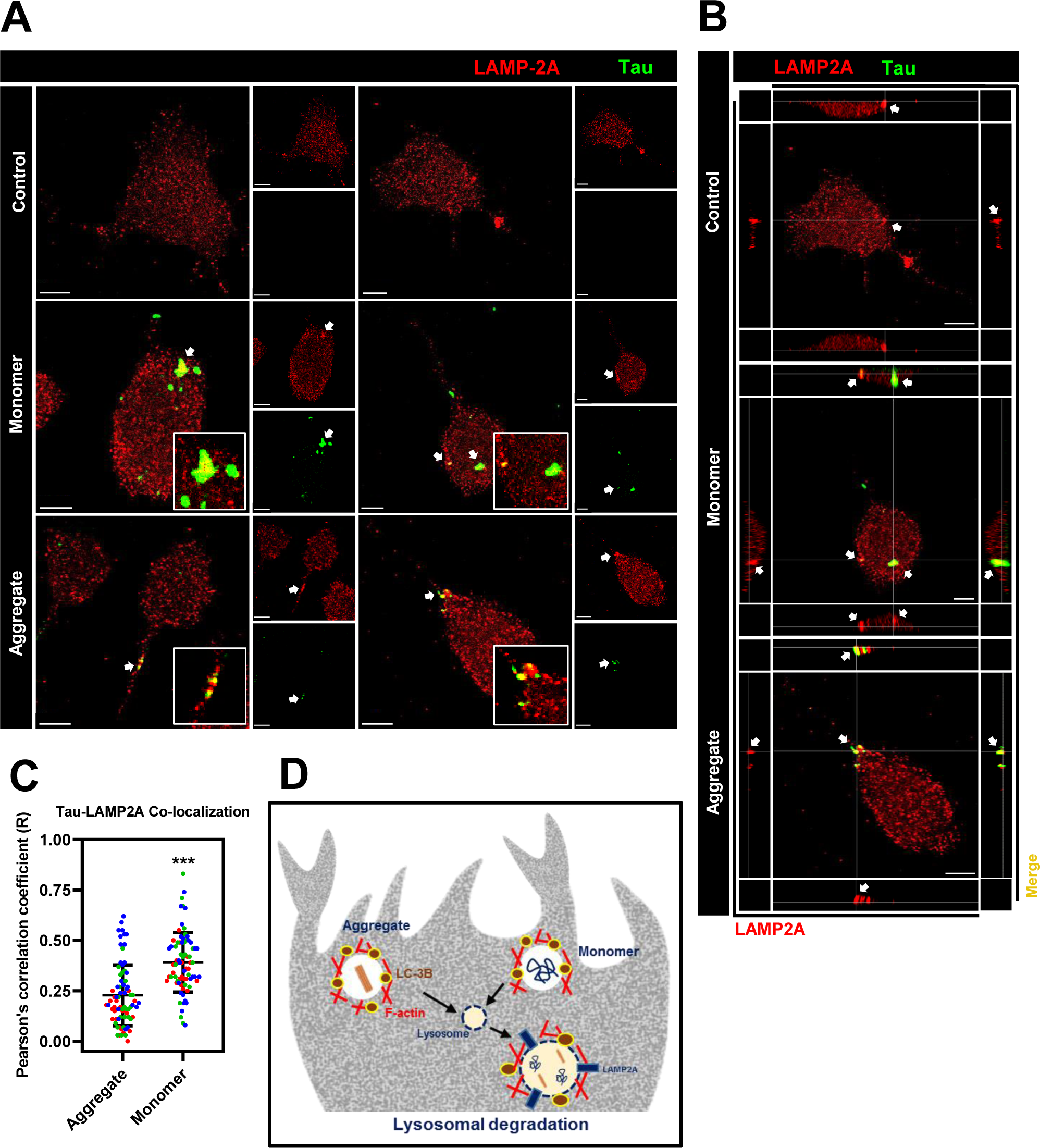
Lysosomal degradation of internalized Tau. (A) Confocal microscope images of N9 microglial cells treated with Alexa Fluor^647^-labeled monomeric and aggregated Tau for 24 h are stained with LAMP2A. White arrows indicate Tau-LAMP2A colocalization. The scale bar represents 5 µm. (B) Orthogonal representation of Tau colocalization with LAMP2A. The left and bottom axes show LAMP2A intensity, whereas the right and top axes represent colocalization between LAMP2A and Tau for better visualization. (C) Pearson’s co-efficient of colocalization plotted for Tau and LAMP2A that shows increased colocalization for Tau monomers compared to aggregates (each color code represents a biological repeat) (*n* = 60). The data is statistically significant with *P* value <0.05 calculated through an unpaired Student’s *t*-test. (D) Diagrammatic representation of lysosomal degradation of extracellular Tau upon LC3- associated endocytosis. *** *P* <0.001, ** *P* <0.01, * *P* <0.05, ^ns^ *P* ≥0.05. LAMP2A: Lysosome-associated membrane protein-2A; LC3: Microtubule-associated protein 1 light chain 3.

### Chloroquine impairs autophagosome-lysosomal fusion and promotes accumulation of autophagosome in microglia

Chloroquine inhibits autophagy by preventing autophagosomal fusion with lysosomes.(^55, 78, 79^) In this study, we used 30 µM chloroquine as a pre-treatment to determine the role of LC3- associated endosomes in Tau degradation by lysosomal pathway. Following extracellular Tau exposure, LC3 distribution in microglial cells was observed after 3 h [Figure 6A]. Membrane association of LC3-II in the Tau-treated groups showed active participation in LANDO (indicated by white arrows). In the chloroquine-treated groups, LC3-associated endosomes accumulated in the perinuclear region of the cells. The orthogonal representation clearly demonstrates the accumulation of LC3-associated endosomes that highly colocalizes with Alexa Fluor^647^-tagged Tau species in chloroquine-treated groups in comparison with the control group [Figure 6B]. As accumulation was observed, we quantified the size of the LC3-associated endosomes by measuring the two-dimensional area of the puncta in different Tau-treated groups. The puncta sizes ranged from 0.02 to 0.2 µm^2^. In the control group, maximum values (∼20%) are observed at 0.01–0.02 µm^2^ whereas the aggregate-treated group demonstrated a slight reduction (∼18%) in the 0.01–0.02 µm^2^ range. Similarly, in the chloroquine-treated groups, the area percentage was further reduced (∼12–14%) in this size range. The percentage area increased at 0.2 µm^2^ in the Tau-treated groups which is further upregulated by chloroquine treatment (indicated by red arrow). This clearly demonstrates that the puncta size increased in the chloroquine-treated groups that shifted toward 0.2 µm^2^ [Figure 6C]. The accumulation of LC3- associated endosomes is also evidenced with the increase in the area percentage of ≥0.5 µm^2^ that are calculated from the accumulated bodies at the perinuclear site [Figure 6C]. We calculated the number of puncta per cell in the individual groups compared with that in the control group. The number of puncta per cell was significantly reduced in the chloroquine-treated groups compared to that in the Tau and control groups [Figure 6D]. These results suggested impaired lysosomal fusion and LC3-associated endosomal accumulation in microglia [Figure 6E].

**Figure 6:**
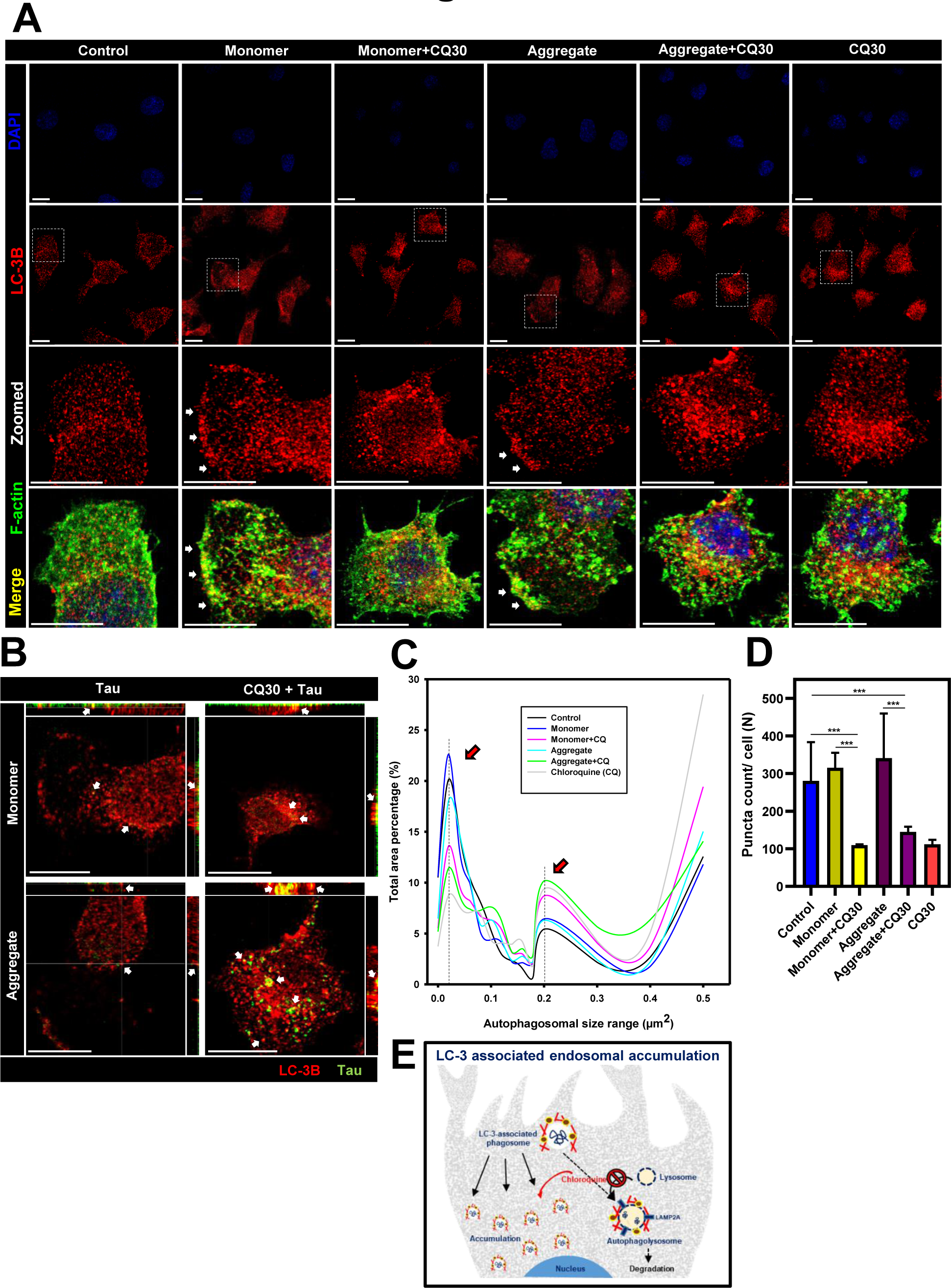
Chloroquine inhibits lysosomal fusion and promotes autophagosomal accumulation. (A) Confocal microscope images of N9 microglial cells treated with monomeric (1 µmol/L) and aggregated Tau (1 µmol/L) for 3 h in the presence or absence of chloroquine (30 µmol/L). The cells are stained for LC3B and DAPI. The scale bar represents 10 µm. (B) Orthogonal representation showing autophagosomal accumulation colocalizing with Alexa Fluor^647^-labeled monomeric and aggregated Tau (white arrows show Tau-LC3 colocalization). (C) Quantification of autophagosome size that ranges from 0.02 to 0.5 µm^2^ by ImageJ software. The x-axis shows the range of autophagosomal size (µm^2^), and y-axis shows the percentage of total area of different size ranges of autophagosomes. The red arrow shows an increase in the area range of autophagosomes upon Tau and chloroquine-treated groups (*n* = 12). (D) Quantification of the number of LC3 puncta per cell in Tau-treated groups as compared to chloroquine and the control group (*n* = 12). The data is statistically significant with a *P* value <0.05 calculated using one-way ANOVA and the statistical significance of the multiple groups has been calculated by Tukey– Kramer’s post hoc analysis for multiple comparisons at 5% level of significance (*** *P* <0.001, ** *P* <0.01, * *P* <0.05, ^ns^ *P* ≥0.05). (E) Schematic representation of LC3-associated autophagosomal accumulation. ANOVA: Analysis of variance; CQ30: Chloroquine (30 µmol/L); DAPI: 4′,6-Diamidino-2-phenylindole; LC3: Microtubule-associated protein 1 light chain 3.

### Chloroquine inhibits lysosomal degradation leading to Tau accumulation in microglia

The degradation of internalized Tau by LANDO was studied by impairing lysosomal fusion in microglial cells following chloroquine exposure and observed under Leica Stellaris 5 confocal microscope. The accumulation of Tau monomers and aggregates in the chloroquine-treated groups was compared with that in the Tau-treated and control groups [Figure 7A]. The mean intensity values were quantified for internalized Tau from individual groups and showed a significant increase in the chloroquine-treated groups, indicating impaired Tau degradation by lysosomes. Tau aggregates were better degraded by lysosomes, which showed a maximum increase in the mean intensity values compared to the monomers [Figure 7B]. These data indicate the lysosomal degradation of Tau monomers and aggregates internalized by LANDO [Figure 7C].

**Figure 7:**
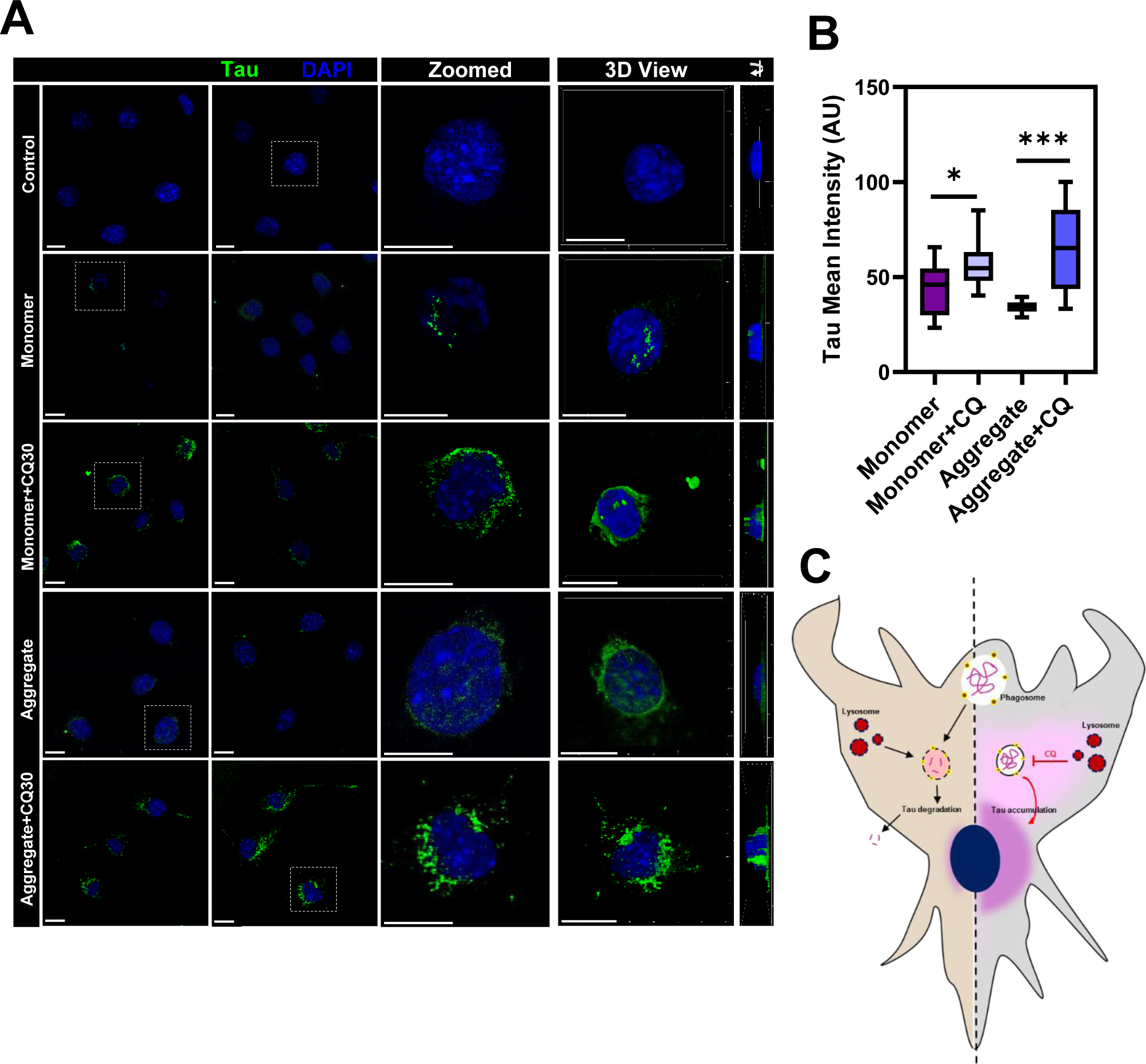
Chloroquine promotes Tau accumulation in microglia. (A) Confocal microscope images of N9 microglial cells treated with Alexa Fluor^647^-labeled monomeric and aggregated Tau for 3 h in the presence and absence of chloroquine (30 μM). Chloroquine is provided 30 min prior to Tau exposure. Chloroquine-exposed cells demonstrating Tau accumulation in monomer and aggregate treated cells as compared to control groups. Scale bar represents 10 µm. (B) Quantification of Tau mean intensity in Tau-treated groups compared to chloroquine-treated groups (*n* = 12). The data is statistically significant with a *P* value <0.05 calculated using one-way ANOVA, and the statistical significance of the multiple groups has been calculated using Tukey–Kramer’s post hoc analysis for multiple comparisons at 5% level of significance (*** *P* <0.001, ** *P* <0.01, * *P* <0.05, ^ns^ *P* ≥0.05). (C) Schematic representation of Tau accumulation in microglia upon autophagosomal inhibition. ANOVA: Analysis of variance; CQ: Chloroquine.

## Discussion

Neuroinflammatory conditions lead to the upregulation of Iba-1 protein levels by migratory microglia in the brain.(^80, 81^) In addition to classical M1 and M2 polarized microglial activation, several other states occur depending on external stimuli and their microenvironments, such as interleukin (IL)-33-responsive microglia, interferon response microglia, proliferative region-associated microglia, lipid-droplet-accumulating microglia, and activated response microglia (ARMs).(^82^) The ARM state is observed in many neurodegenerative diseases such as AD and is also known as disease-associated microglia (DAM). Extracellular signals secreted from degenerative neurons, dead cells, and misfolded protein aggregates are believed to play key roles in microglial activation.(^83^) Several receptors are even reported to be involved in promoting the DAM phenotype of microglia such as the triggering receptor expressed on myeloid cell 2 (TREM-2) receptor and apolipoprotein E (ApoE) receptor which plays a crucial role in sensing the extracellular protein aggregates such as Tau, Aβ, and α-synuclein.(^84–87^) DAM is associated with enhanced lipid metabolism, increased phagocytic activity, lysosomal protease synthesis for active degradation, secretion of neuroprotective molecules, and restoration of homeostasis.(^84^) Extracellular Tau species promote microglial activation for its active internalization.(^25, 36, 88, 89^) In our study, extracellular monomer and aggregate Tau treatments activated microglia that underwent actin polymerization and membrane ruffling at the leading edge, which highly colocalized with the Iba-1 marker. The enhanced Iba-1 expression confirms the activated state of microglia.(^71^) The three-dimensional representation visualized the membrane ruffles in the treatment groups, denoting microglial endocytosis,(^70^) which is comparable to our previous studies.(25, 36, 90)

Autophagy plays a significant role in several neurodegenerative diseases, such as Parkinson’s disease, Huntington’s disease, and AD through the clearance of intracellular protein aggregates, such as α-synuclein and Tau.(^40, 91^) Autophagy inhibitors such as 3-methyl adenine (3MA) and bafilomycin A1 (Baf) delay cytosolic Tau and α-synuclein clearance, whereas rapamycin, an autophagy inducer, promotes clearance of these proteins from the cytosolic space.(92–94) *Drosophila* expressing R406W Tau leads to the formation of abnormal or absence of eyes. Rapamycin alleviated Tau-mediated toxicity and reduced the number of the “no eyes” populations in the rapamycin-treated group.(^92^) Microglia is reported to prevent neurodegeneration by engulfing neuron-released α-synuclein through a selective autophagy mechanism called synucleinphagy.(^95^) Similar to LAP, LANDO is another non-canonical pathway for the ingestion of toxic extracellular materials and misfolded protein aggregates, where the endosomes associate with the autophagy marker LC3 for internalization.(^96, 97^) However, the exact molecular factors underlying the activation of LANDO need to be further explored. LAP is triggered by the activation of several cell surface receptors, such as TLRs (1/2, 2/6, and 4), T-cell immunoglobulin mucin protein 4 (TIM4), and Fc receptor (FcR). Contrarily, LAP and canonical autophagy activation have independent mechanisms. Autophagy depends on activation kinases, such as mammalian target of rapamycin complex 1 (mTORC1) and adenosine monophosphate-activated protein kinase (AMPK), which are mediated by the pre-initiation complex. Conversely, LAP activation is based on the triggering of surface receptors and is independent of the pre-initiation complex.(^98, 99^) LAP has a significant role in efficient clearance of dead cells(^100^) engulfing viable bacteria such as *Listeria monocytogenes* by macrophages,(^101^) and antifungal immunity against *Saccharomyces cerevisiae* and *Aspergillus fumigatus*.(^51, 99^) In AD, recent studies have elucidated the role of LANDO in the clearance of Aβ plaques.(^52^) Autophagy regulators ATG5 and Rubicon-deficient mice exacerbated Aβ accumulation in the cortex and hippocampus with no significant changes in the Focal adhesion kinase family interacting protein of 200 kD (FIP200) deficient mice, which denoted a non-canonical autophagy mechanism. Moreover, LC3-containing vesicles co-localized with clathrin and Ras-associated binding protein-5 (Rab5), irrespective of the inhibition of phagocytosis by latrunculin A. Additionally, LANDO plays a key role in TREM-2 receptor recycling from receptor-mediated endocytosis of Aβ, regulates Tau hyperphosphorylation, and protects against Aβ-induced microglial activation, neuronal loss, and behavioral and memory impairment in murine AD.(^52^) In previous studies, we showed that Tau endocytosis and vesicular trafficking of extracellular oligomers and aggregates are mediated by surface receptors, such as P2Y12R and CX3CR1, which fuse with lysosomes for degradation.(^25, 26, 37^) Tau internalization by clathrin-mediated endocytic pathways is a well-studied mechanism that promotes vesicular trafficking of Tau through early and late endosomes for lysosomal degradation.(^37, 75^) In the present study, we analyzed the role of LANDO in the internalization and degradation of extracellular Tau by microglial cells. We demonstrated that microglial cells internalize extracellular Tau that colocalized with LC3 containing endosomes showing LC3-associated Tau endocytosis followed by lysosomal degradation.

In the early onset of AD, autophagy is induced for the active clearance of Aβ and Tau by forming autophagosomes in neuronal cells.(^102^) Later, autophagic pathology and accumulation occurred because of impaired lysosomal degradation.(^53^) In the later stages of AD, rapamycin causes autophagic stress in cells, probably owing to the reduced lysosomal degradative potential.(^103^) Our study reveals the LAP-associated lysosomal degradation of extracellular Tau through active colocalization with LAMP2A could possibly have a beneficial role in the early stages of disease progression. Furthermore, we demonstrated lysosomal degradation of LC3- associated endosomes by the autophagy inhibitor chloroquine. Chloroquine was previously considered a lysosomal inhibitor that prevents lysosomal acidification.(^104^) Later, Mauthe *et al*, reported the mechanism of action of chloroquine in lysosomal degradation. Chloroquine inhibits autophagosomal bulk degradation without changing lysosomal acidity by impairing autophagosome-lysosomal fusion.(^55, 78, 79^) It is also inferred that aggregated Tau could be better cleared by lysosomes than monomers, which is evident from significant Tau accumulation in chloroquine-treated groups. Although several mechanisms of Tau clearance by microglia have been reported in the early stages of AD pathology, we have shown that LANDO is a novel method of Tau clearance by microglial cells that can limit Tau spreading and promote lysosomal degradation under tauopathies [Figure 8].

**Figure 8:**
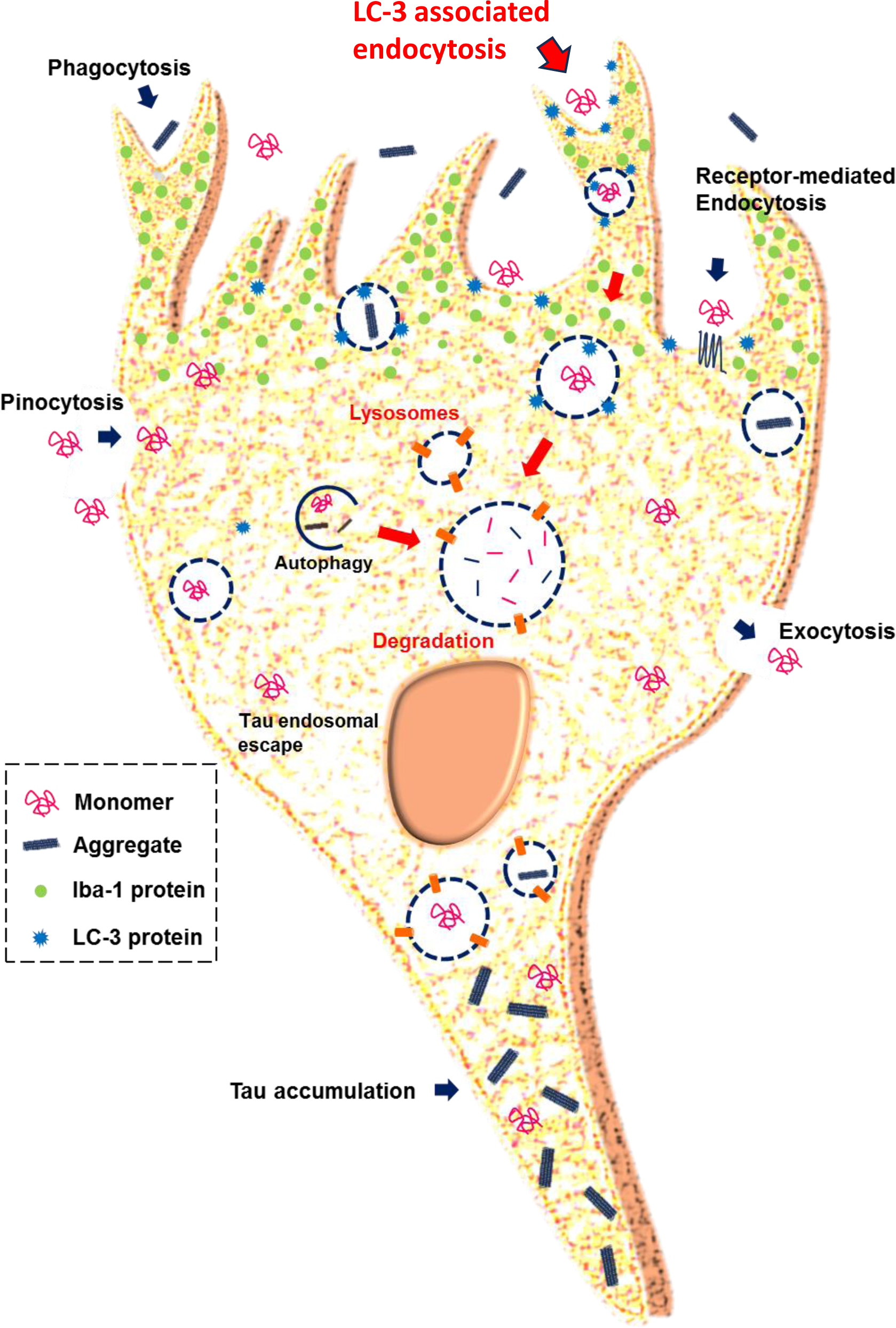
Summary of Tau transmission and LC3 associated endocytosis in microglia. Tau is internalized by microglia through several mechanisms that include micropinocytosis, HSPG- dependent pathways, receptor-mediated endocytosis, and phagocytosis. Internalized Tau is accumulated in the cytosol of microglial cells. LC3-associated Tau endocytosis could be an efficient mechanism of Tau clearance to be further studied followed by lysosomal degradation. HSPG: Heparan sulfate proteoglycan; LC3: Microtubule-associated protein 1 light chain 3.

In conclusion, extracellular Tau activates microglial cells. Activated microglia undergo LANDO for extracellular Tau aggregate clearance via active internalization and lysosomal degradation, suggesting a molecular mechanism that could be further studied in regulating tauopathy and neurodegeneration.

## Acknowledgments

The authors thank the Department of Biotechnology for the fellowship. We thank Rashmi Das and Smita Desale for their valuable comments and suggestions for performing the cell biology experiments. We are grateful to the members of Chinnathambi’s lab for their scientific discussions, helpful suggestions, and critical reading of the manuscript.

## References

1. Arcuri C, Mecca C, Bianchi R, Giambanco I, Donato R. The pathophysiological role of microglia in dynamic surveillance, phagocytosis and structural remodeling of the developing CNS. Frontiers in Molecular Neuroscience. 2017;10:191. doi: 10.3389/fnmol.2017.00191.

2. Nimmerjahn A, Kirchhoff F, Helmchen F. Resting microglial cells are highly dynamic surveillants of brain parenchyma in vivo. Science. 2005;308(5726):1314-8. doi: 10.1126/science.1110647.

3. Perea JR, Bolós M, Avila J. Microglia in Alzheimer’s disease in the context of tau pathology. Biomolecules. 2020;10(10):1439. doi: 10.3390/biom10101439.

4. Davalos D, Grutzendler J, Yang G, Kim JV, Zuo Y, Jung S, et al. ATP mediates rapid microglial response to local brain injury in vivo. Nature neuroscience. 2005;8(6):752–8. doi: 10.1038/nn1472.

5. Hickman SE, Kingery ND, Ohsumi TK, Borowsky ML, Wang L-c, Means TK, et al. The microglial sensome revealed by direct RNA sequencing. Nature neuroscience. 2013;16(12):1896–905. doi: 10.1038/nn.3554.

6. Haynes SE, Hollopeter G, Yang G, Kurpius D, Dailey ME, Gan W-B, et al. The P2Y12 receptor regulates microglial activation by extracellular nucleotides. Nature neuroscience. 2006;9(12):1512–9. doi: 10.1038/nn1805.

7. Madry C, Kyrargyri V, Arancibia-Cárcamo IL, Jolivet R, Kohsaka S, Bryan RM, et al. Microglial ramification, surveillance, and interleukin-1β release are regulated by the two-pore domain K+ channel THIK-1. Neuron. 2018;97(2):299–312. e6. doi: 10.1016/j.neuron.2017.12.002.

8. Noda M, Nakanishi H, Nabekura J, Akaike N. AMPA–kainate subtypes of glutamate receptor in rat cerebral microglia. Journal of Neuroscience. 2000;20(1):251–8. doi: 10.1523/JNEUROSCI.20-01-00251.2000.

9. Chinnathambi S, Gorantla NV. Implications of Valosin-containing Protein in Promoting Autophagy to Prevent Tau Aggregation. Neuroscience. 2021;476:125–34. doi: 10.1016/j.neuroscience.2021.09.003.

10. Binder LI, Frankfurter A, Rebhun LI. The distribution of tau in the mammalian central nervous system. Journal of Cell Biology. 1985;101(4):1371–8. doi: 10.1083/jcb.101.4.1371.

11. Conde C, Cáceres A. Microtubule assembly, organization and dynamics in axons and dendrites. Nature Reviews Neuroscience. 2009;10(5):319. doi: 10.1038/nrn2631.

12. Sonawane SK, Chinnathambi S. Prion-like propagation of post-translationally modified tau in Alzheimer’s disease: a hypothesis. Journal of Molecular Neuroscience. 2018;65(4):480–90. doi: 10.1007/s12031-018-1111-5.

13. Gong C-X, Liu F, Grundke-Iqbal I, Iqbal K. Post-translational modifications of tau protein in Alzheimer’s disease. Journal of neural transmission. 2005;112(6):813–38. doi: 10.1007/s00702-004-0221-0.

14. Martin L, Latypova X, Terro F. Post-translational modifications of tau protein: implications for Alzheimer’s disease. Neurochemistry international. 2011;58(4):458–71. doi: 10.1016/j.neuint.2010.12.023.

15. Das R, Chinnathambi S. Microglial priming of antigen presentation and adaptive stimulation in Alzheimer’s disease. Cellular and Molecular Life Sciences. 2019;76(19):3681–94. doi: 10.1007/s00018-019-03132-2.

16. Kanmert D, Cantlon A, Muratore CR, Jin M, O’Malley TT, Lee G, et al. C-terminally truncated forms of tau, but not full-length tau or its C-terminal fragments, are released from neurons independently of cell death. Journal of Neuroscience. 2015;35(30):10851–65. doi: 10.1523/JNEUROSCI.0387-15.2015.

17. Pérez M, Avila J, Hernández F. Propagation of tau via extracellular vesicles. Frontiers in Neuroscience. 2019;13:698. doi: 10.3389/fnins.2019.00698.

18. Pooler AM, Phillips EC, Lau DH, Noble W, Hanger DP. Physiological release of endogenous tau is stimulated by neuronal activity. EMBO reports. 2013;14(4):389–94. doi: 10.1038/embor.2013.15.

19. Yamada K, Holth JK, Liao F, Stewart FR, Mahan TE, Jiang H, et al. Neuronal activity regulates extracellular tau in vivo. Journal of Experimental Medicine. 2014;211(3):387–93. doi: 10.1084/jem.20131685.

20. Chinnathambi S. α-Linolenic Acid Vesicles-Mediated Tau Internalization in Microglia. Lipid Signalling: Methods and Protocols: Springer; 2024. p. 117–28. doi: 10.1007/978-1-0716-3902-3_11.

21. Perea JR, López E, Díez-Ballesteros JC, Ávila J, Hernández F, Bolós M. Extracellular monomeric tau is internalized by astrocytes. Frontiers in neuroscience. 2019;13:442. doi: 10.3389/fnins.2019.00442.

22. Kolay S, Vega AR, Dodd DA, Perez VA, Kashmer OM, White CL, et al. The dual fates of exogenous tau seeds: Lysosomal clearance versus cytoplasmic amplification. Journal of Biological Chemistry. 2022;298:102014. doi: 10.1016/j.jbc.2022.102014.

23. Evans LD, Wassmer T, Fraser G, Smith J, Perkinton M, Billinton A, et al. Extracellular monomeric and aggregated tau efficiently enter human neurons through overlapping but distinct pathways. Cell reports. 2018;22(13):3612–24. doi: 10.1016/j.celrep.2018.03.021.

24. Bolós M, Llorens-Martín M, Perea JR, Jurado-Arjona J, Rábano A, Hernández F, et al. Absence of CX3CR1 impairs the internalization of Tau by microglia. Molecular neurodegeneration. 2017;12(1):59. doi: 10.1186/s13024-017-0200-1.

25. Chidambaram H, Das R, Chinnathambi S. G-Protein coupled Purinergic P2Y12 receptor interacts and internalizes TauRD-mediated by membrane-associated actin cytoskeleton remodelling in microglia. European Journal of Cell Biology. 2022:151201. doi: 10.1016/j.ejcb.2022.151201.

26. Das R, Chinnathambi S. Microglial remodeling of actin network by Tau oligomers, via G protein-coupled purinergic receptor, P2Y12R- driven chemotaxis. Traffic. 2021;22(5):153-70. doi: 10.1111/tra.12784.

27. Chinnathambi S, Chidambaram H. G-protein coupled receptors regulates Tauopathy in neurodegeneration. Advances in Protein Chemistry and Structural Biology. 2024;141:467–93. doi: 10.1016/bs.apcsb.2024.04.001.

28. Chidambaram H, Desale SE, Chinnathambi S. Purinergic Receptor P2Y12-Mediated Tau Internalization in Microglia. Tau Protein: Methods and Protocols: Springer; 2024. p. 457–70. doi: 10.1007/978-1-0716-3629-9_25.

29. Chidambaram H, Desale SE, Chinnathambi S. Interaction of Tau with G-Protein-Coupled Purinergic P2Y12 Receptor by Molecular Docking and Molecular Dynamic Simulation. Tau Protein: Methods and Protocols: Springer; 2024. p. 33–54. doi: 10.1007/978-1-0716-3629-9_2.

30. Chidambaram H, Chinnathambi S. G-protein coupled receptors and tau-different roles in Alzheimer’s disease. Neuroscience. 2020;438:198–214. doi: 10.1016/j.neuroscience.2020.04.019.

31. Chinnathambi S, Das R. Microglia degrade Tau oligomers deposit via purinergic P2Y12- associated podosome and filopodia formation and induce chemotaxis. Cell & Bioscience. 2023;13(1):95. doi: 10.1186/s13578-023-01028-0.

32. Chinnathambi S, Das R, Desale SE. Tau aggregates improve the Purinergic receptor P2Y12-associated podosome rearrangements in microglial cells. Biochimica et Biophysica Acta (BBA)-Molecular Cell Research. 2023;1870(5):119477. doi: 10.1016/j.bbamcr.2023.119477.

33. Uribe-Querol E, Rosales C. Control of phagocytosis by microbial pathogens. Frontiers in immunology. 2017;8:1368. doi:10.3389/fimmu.2017.01368.

34. Chidambaram H, Desale SE, Qureshi T, Chinnathambi S. Microglial Uptake of Extracellular Tau by Actin-Mediated Phagocytosis. Neuroprotection: Method and Protocols: Springer; 2024. p. 231–43. doi: 10.1007/978-1-0716-3662-6_16.

35. Chinnathambi S, Desale SE. The crosstalk between extracellular matrix proteins and Tau. Advances in Protein Chemistry and Structural Biology. 2024;141:447–66. doi: 10.1016/bs.apcsb.2024.04.002.

36. Das R, Balmik AA, Chinnathambi S. Phagocytosis of full-length Tau oligomers by Actin-remodeling of activated microglia. Journal of neuroinflammation. 2020;17(1):1–15. doi: 10.1186/s12974-019-1694-y.

37. Desale SE, Chinnathambi S. α–Linolenic acid modulates phagocytosis and endosomal pathways of extracellular Tau in microglia. Cell adhesion & migration. 2021;15(1):84–100. doi: 10.1080/19336918.2021.1898727.

38. Desale SE, Chidambaram H, Qureshi T, Chinnathambi S. Internalization and Endosomal Trafficking of Extracellular Tau in Microglia Improved by α-Linolenic Acid. Neuroprotection: Method and Protocols: Springer; 2024. p. 245–55. doi:10.1007/978-1-0716-3662-6_17.

39. Ravikumar B, Moreau K, Jahreiss L, Puri C, Rubinsztein DC. Plasma membrane contributes to the formation of pre-autophagosomal structures. Nature cell biology. 2010;12(8):747–57. doi: 10.1038/ncb2078.

40. Ravikumar B, Sarkar S, Davies JE, Futter M, Garcia-Arencibia M, Green-Thompson ZW, et al. Regulation of mammalian autophagy in physiology and pathophysiology. Physiological reviews. 2010;90(4):1383–435. doi: 10.1152/physrev.00030.2009.

41. Kabeya Y, Mizushima N, Ueno T, Yamamoto A, Kirisako T, Noda T, et al. LC3, a mammalian homologue of yeast Apg8p, is localized in autophagosome membranes after processing. The EMBO journal. 2000;19(21):5720–8. doi: 10.1093/emboj/19.21.5720.

42. Klionsky DJ, Abdel-Aziz AK, Abdelfatah S, Abdellatif M, Abdoli A, Abel S, et al. Guidelines for the use and interpretation of assays for monitoring autophagy. autophagy. 2021;17(1):1–382. doi: 10.1080/15548627.2020.1797280.

43. Schläfli A, Berezowska S, Adams O, Langer R, Tschan M. Reliable LC3 and p62 autophagy marker detection in formalin fixed paraffin embedded human tissue by immunohistochemistry. European journal of histochemistry: EJH. 2015;59(2). doi: 10.4081/ejh.2015.2481.

44. Kabeya Y, Mizushima N, Yamamoto A, Oshitani-Okamoto S, Ohsumi Y, Yoshimori T. LC3, GABARAP and GATE16 localize to autophagosomal membrane depending on form-II formation. Journal of cell science. 2004;117(13):2805-12. doi: 10.1242/jcs.01131.

45. Bento CF, Renna M, Ghislat G, Puri C, Ashkenazi A, Vicinanza M, et al. Mammalian autophagy: how does it work. Annu Rev Biochem. 2016;85(1):685–713. doi: 10.1146/annurev-biochem-060815-014556.

46. Eskelinen E-L, Saftig P. Autophagy: a lysosomal degradation pathway with a central role in health and disease. Biochimica et Biophysica Acta (BBA)-Molecular Cell Research. 2009;1793(4):664–73. doi: 10.1016/j.bbamcr.2008.07.014.

47. Deen NS, Gong L, Naderer T, Devenish RJ, Kwok T. Analysis of the Relative Contribution of Phagocytosis, LC 3C 3Biochimd Phagocytosis, and Canonical Autophagy During Helicobacter pylori Infection of Macrophages. Helicobacter. 2015;20(6):449-59. doi: 10.1111/hel.12223.

48. Kageyama S, Omori H, Saitoh T, Sone T, Guan J-L, Akira S, et al. The LC3 recruitment mechanism is separate from Atg9L1-dependent membrane formation in the autophagic response against Salmonella. Molecular biology of the cell. 2011;22(13):2290–300. doi: 10.1091/mbc.E10-11-0893.

49. Oikonomou V, Renga G, De Luca A, Borghi M, Pariano M, Puccetti M, et al. Autophagy and LAP in the fight against fungal infections: Regulation and therapeutics. Mediators of inflammation. 2018;2018. doi: 10.1155/2018/6195958.

50. Sprenkeler EG, Gresnigt MS, van de Veerdonk FL. LC3LC3Puccetti M, et al. Autophagy and LAP in the fight against fungal infections: Regulation and therapeutics. Mediators of inflamma16;18(9):1208-16. doi: 10.1111/cmi.12616.

51. Sanjuan MA, Dillon CP, Tait SW, Moshiach S, Dorsey F, Connell S, et al. Toll-like receptor signalling in macrophages links the autophagy pathway to phagocytosis. Nature. 2007;450(7173):1253-7. doi: 10.1038/nature06421.

52. Heckmann BL, Teubner BJ, Tummers B, Boada-Romero E, Harris L, Yang M, et al. LC3-associated endocytosis facilitates β-amyloid clearance and mitigates neurodegeneration in murine Alzheimer’s disease. Cell. 2019;178(3):536–51. e14. doi: 10.1016/j.cell.2019.05.056.

53. Piras A, Collin L, Grüninger F, Graff C, Rönnbäck A. Autophagic and lysosomal defects in human tauopathies: analysis of post-mortem brain from patients with familial Alzheimer disease, corticobasal degeneration and progressive supranuclear palsy. Acta neuropathologica communications. 2016;4(1):1–13. doi: 10.1186/s40478-016-0292-9.

54. Frosch T, Schmitt M, Bringmann G, Kiefer W, Popp J. Structural analysis of the anti-malaria active agent chloroquine under physiological conditions. The Journal of Physical Chemistry B. 2007;111(7):1815–22. doi: 10.1021/jp065136j.

55. Mauthe M, Orhon I, Rocchi C, Zhou X, Luhr M, Hijlkema K-J, et al. Chloroquine inhibits autophagic flux by decreasing autophagosome-lysosome fusion. Autophagy. 2018;14(8):1435–55. doi: 10.1080/15548627.2018.1474314.

56. Chidambaram H, Chinnathambi S. Role of cysteines in accelerating Tau filament formation. Journal of Biomolecular Structure and Dynamics. 2020:1–10. doi: 10.1080/07391102.2020.1856720.

57. Gorantla NV, Das R, Chidambaram H, Dubey T, Mulani FA, Thulasiram HV, et al. Basic Limonoid modulates Chaperone-mediated Proteostasis and dissolve Tau fibrils. Scientific Reports. 2020;10(1):4023. doi: 10.1038/s41598-020-60773-1.

58. Sonawane SK, Chidambaram H, Boral D, Gorantla NV, Balmik AA, Dangi A, et al. EGCG impedes human Tau aggregation and interacts with Tau. Scientific reports. 2020;10(1):12579. doi: 10.1038/s41598-020-69429-6.

59. Dangi A, Qureshi T, Chinnathambi S, Marelli UK. Macrocyclic peptides derived from AcPHF6* and AcPHF6 to selectively modulate the Tau aggregation. Bioorganic Chemistry. 2024;151:107625. doi: 10.1016/j.bioorg.2024.107625.

60. Desale SE, Chidambaram H, Chinnathambi S. Biochemical and Biophysical Characterization of Tau and α-Linolenic Acid Vesicles In Vitro. Tau Protein: Methods and Protocols: Springer; 2024. p. 193–203. doi: 10.1007/978-1-0716-3629-9_11.

61. Dubey T, Kushwaha P, Thulasiram H, Chandrashekar M, Chinnathambi S. Bacopa monnieri reduces Tau aggregation and Tau-mediated toxicity in cells. International journal of biological macromolecules. 2023;234:123171. doi: 10.1016/j.ijbiomac.2023.123171.

62. Balmik AA, Chidambaram H, Dangi A, Marelli UK, Chinnathambi S. HDAC6 ZnF UBP as the modifier of tau structure and function. Biochemistry. 2020;59(48):4546–62. doi: 10.1021/acs.biochem.0c00585.

63. Friedhoff P, Schneider A, Mandelkow E-M, Mandelkow E. Rapid assembly of Alzheimer-like paired helical filaments from microtubule-associated protein tau monitored by fluorescence in solution. Biochemistry. 1998;37(28):10223–30. doi: 10.1021/bi980537d.

64. Gorantla NV, Das R, Chidambaram H, Dubey T, Mulani FA, Thulasiram HV, et al. Basic Limonoid modulates Chaperone-mediated Proteostasis and dissolve Tau fibrils. Scientific reports. 2020;10(1):1–19. doi: 10.1038/s41598-020-60773-1.

65. Kampers T, Friedhoff P, Biernat J, Mandelkow E-M, Mandelkow E. RNA stimulates aggregation of microtubule-associated protein tau into Alzheimer-like paired helical filaments. FEBS letters. 1996;399(3):344–9. doi:10.1016/s0014-5793(96)01386-5.

66. Sonawane SK, Balmik AA, Boral D, Ramasamy S, Chinnathambi S. Baicalein suppresses Repeat Tau fibrillization by sequestering oligomers. Archives of biochemistry and biophysics. 2019;675:108119. doi: 10.1016/j.abb.2019.108119.

67. Balmik AA, Das R, Dangi A, Gorantla NV, Marelli UK, Chinnathambi S. Melatonin interacts with repeat domain of Tau to mediate disaggregation of paired helical filaments. Biochimica et Biophysica Acta (BBA)-General Subjects. 2020;1864(3):129467. doi: 10.1016/j.bbagen.2019.129467.

68. Wobst HJ, Sharma A, Diamond MI, Wanker EE, Bieschke J. The green tea polyphenol (−)-epigallocatechin gallate prevents the aggregation of tau protein into toxic oligomers at substoichiometric ratios. FEBS letters. 2015;589(1):77–83. doi: 10.1016/j.febslet.2014.11.026.

69. Jin M, Shiwaku H, Tanaka H, Obita T, Ohuchi S, Yoshioka Y, et al. Tau activates microglia via the PQBP1-cGAS-STING pathway to promote brain inflammation. Nature communications. 2021;12(1):1–22. doi: 10.1038/s41467-021-26851-2.

70. Ohsawa K, Imai Y, Kanazawa H, Sasaki Y, Kohsaka S. Involvement of Iba1 in membrane ruffling and phagocytosis of macrophages/microglia. Journal of cell science. 2000;113(17):3073–84. doi: 10.1242/jcs.113.17.3073.

71. Ito D, Imai Y, Ohsawa K, Nakajima K, Fukuuchi Y, Kohsaka S. Microglia-specific localisation of a novel calcium binding protein, Iba1. Molecular brain research. 1998;57(1):1-9. doi: 10.1016/s0169-328x(98)00040-0.

72. Perez-Pouchoulen M, VanRyzin JW, McCarthy MM. Morphological and phagocytic profile of microglia in the developing rat cerebellum. eneuro. 2015;2(4). doi: 10.1523/ENEURO.0036-15.2015.

73. Swanson JA. Shaping cups into phagosomes and macropinosomes. Nature reviews Molecular cell biology. 2008;9(8):639–49. doi: 10.1038/nrm2447.

74. Chidambaram H, Das R, Chinnathambi S. Interaction of Tau with the chemokine receptor, CX3CR1 and its effect on microglial activation, migration and proliferation. Cell & bioscience. 2020;10(1):1-9. doi: 10.1186/s13578-020-00474-4.

75. Zhao J, Wu H, Tang X-q. Tau internalization: A complex step in tau propagation. Ageing Research Reviews. 2021;67:101272. doi: 10.1016/j.arr.2021.101272.

76. Mizushima N. Physiological functions of autophagy. Autophagy in Infection and Immunity. 2009:71–84. doi: 10.1007/978-3-642-00302-8_3.

77. Reinstein E, Ciechanover A. Narrative review: protein degradation and human diseases: the ubiquitin connection. Annals of internal medicine. 2006;145(9):676–84. doi: 10.7326/0003-4819-145-9-200611070-00010.

78. Bik E, Mateuszuk L, Orleanska J, Baranska M, Chlopicki S, Majzner K. Chloroquine-induced accumulation of autophagosomes and lipids in the endothelium. International Journal of Molecular Sciences. 2021;22(5):2401. doi: 10.3390/ijms22052401.

79. Neill T, Chen CG, Buraschi S, Iozzo RV. Catabolic degradation of endothelial VEGFA via autophagy. Journal of Biological Chemistry. 2020;295(18):6064–79. doi: 10.1074/jbc.RA120.012593.

80. Seminotti B, Zanatta Â, Ribeiro RT, da Rosa MS, Wyse AT, Leipnitz G, et al. Disruption of brain redox homeostasis, microglia activation and neuronal damage induced by intracerebroventricular administration of S-adenosylmethionine to developing rats. Molecular Neurobiology. 2019;56(4):2760–73. doi: 10.1007/s12035-018-1275-6.

81. Wang L, Jiang Q, Chu J, Lin L, Li X-G, Chai G-S, et al. Expression of Tau40 induces activation of cultured rat microglial cells. PLoS One. 2013;8(10):e76057. doi: 10.1371/journal.pone.0076057.

82. Van Acker ZP, Perdok A, Bretou M, Annaert W. The microglial lysosomal system in Alzheimer’s disease: Guardian against proteinopathy. Ageing Research Reviews. 2021;71:101444. doi: 10.1016/j.arr.2021.101444.

83. García-Revilla J, Alonso-Bellido IM, Burguillos MA, Herrera AJ, Espinosa-Oliva AM, Ruiz R, et al. Reformulating pro-oxidant microglia in neurodegeneration. Journal of Clinical Medicine. 2019;8(10):1719. doi: 10.3390/jcm8101719.

84. Keren-Shaul H, Spinrad A, Weiner A, Matcovitch-Natan O, Dvir-Szternfeld R, Ulland TK, et al. A unique microglia type associated with restricting development of Alzheimer’s disease. Cell. 2017;169(7):1276–90. e17. doi: 10.1016/j.cell.2017.05.018.

85. Weigand AJ, Thomas KR, Bangen KJ, Eglit GM, Delano-Wood L, Gilbert PE, et al. APOE interacts with tau PET to influence memory independently of amyloid PET in older adults without dementia. Alzheimer’s & Dementia. 2021;17(1):61–9. doi: 10.1002/alz.12173.

86. Paslawski W, Zareba-Paslawska J, Zhang X, Hölzl K, Wadensten H, Shariatgorji M, et al. α-synuclein− lipoprotein interactions and elevated ApoE level in cerebrospinal fluid from Parkinson’s disease patients. Proceedings of the National Academy of Sciences. 2019;116(30):15226–35. doi: 10.1073/pnas.1821409116.

87. Zhao Y, Wu X, Li X, Jiang L-L, Gui X, Liu Y, et al. TREM2 is a receptor for β-amyloid that mediates microglial function. Neuron. 2018;97(5):1023–31. e7. doi: 10.1016/j.neuron.2018.01.031.

88. Desale SE, Chinnathambi S. α-Linolenic acid induces clearance of Tau seeds via Actin-remodeling in Microglia. Molecular Biomedicine. 2021;2(1):1–14. 10.1186/s43556-021-00028-1.

89. Desale SE, Chidambaram H, Chinnathambi S. α-Linolenic Acid Induces Microglial Activation and Extracellular Tau Internalization. Tau Protein: Methods and Protocols: Springer; 2024. p. 471–81. doi: 10.1007/978-1-0716-3629-9_26.

90. Desale SE, Chinnathambi S. α-Linolenic Acid Modulates Phagocytosis of Extracellular Tau and Induces Microglial Migration. bioRxiv 2020. doi: 10.1101/2020.04.15.042143.

91. Ravikumar B, Duden R, Rubinsztein DC. Aggregate-prone proteins with polyglutamine and polyalanine expansions are degraded by autophagy. Human molecular genetics. 2002;11(9):1107–17. doi: 10.1093/hmg/11.9.1107.

92. Berger Z, Ravikumar B, Menzies FM, Oroz LG, Underwood BR, Pangalos MN, et al. Rapamycin alleviates toxicity of different aggregate-prone proteins. Human molecular genetics. 2006;15(3):433–42. doi: 10.1093/hmg/ddi458.

93. Noda T, Ohsumi Y. Tor, a phosphatidylinositol kinase homologue, controls autophagy in yeast. Journal of Biological Chemistry. 1998;273(7):3963–6. doi: 10.1074/jbc.273.7.3963.

94. Webb JL, Ravikumar B, Rubinsztein DC. Microtubule disruption inhibits autophagosome-lysosome fusion: implications for studying the roles of aggresomes in polyglutamine diseases. The international journal of biochemistry & cell biology. 2004;36(12):2541–50. doi: 10.1016/j.biocel.2004.02.003.

95. Choi I, Zhang Y, Seegobin SP, Pruvost M, Wang Q, Purtell K, et al. Microglia clear neuron-released α-synuclein via selective autophagy and prevent neurodegeneration. Nature communications. 2020;11(1):1386. doi: 10.1038/s41467-020-15119-w.

96. Herb M, Gluschko A, Schramm M, editors. LC3-associated phagocytosis-The highway to hell for phagocytosed microbes. Seminars in cell & developmental biology; 2020: Elsevier. doi: 10.1016/j.semcdb.2019.04.016.

97. Lai S-c, Devenish RJ. LC3-associated phagocytosis (LAP): connections with host autophagy. Cells. 2012;1(3):396–408. doi: 10.3390/cells1030396.

98. Hurley JH, Young LN. Mechanisms of autophagy initiation. Annual review of biochemistry. 2017;86:225. doi: 10.1146/annurev-biochem-061516-044820.

99. Martinez J, Malireddi R, Lu Q, Cunha LD, Pelletier S, Gingras S, et al. Molecular characterization of LC3-associated phagocytosis reveals distinct roles for Rubicon, NOX2 and autophagy proteins. Nature cell biology. 2015;17(7):893-906. doi: 10.1038/ncb3192.

100. Martinez J, Almendinger J, Oberst A, Ness R, Dillon CP, Fitzgerald P, et al. Microtubule-associated protein 1 light chain 3 alpha (LC3)-associated phagocytosis is required for the efficient clearance of dead cells. Proceedings of the National Academy of Sciences. 2011;108(42):17396–401. doi: 10.1073/pnas.1113421108.

101. Herb M, Gluschko A, Schramm M. LC3-associated phagocytosis initiated by integrin ITGAM-ITGB2/Mac-1 enhances immunity to Listeria monocytogenes. Autophagy. 2018;14(8):1462–4. doi: 10.1080/15548627.2018.1475816.

102. Bordi M, Berg MJ, Mohan PS, Peterhoff CM, Alldred MJ, Che S, et al. Autophagy flux in CA1 neurons of Alzheimer hippocampus: Increased induction overburdens failing lysosomes to propel neuritic dystrophy. Autophagy. 2016;12(12):2467–83. doi: 10.1080/15548627.2016.1239003.

103. Carosi JM, Sargeant TJ. Rapamycin and Alzheimer disease: a hypothesis for the effective use of rapamycin for treatment of neurodegenerative disease. Autophagy. 2023:1–5. doi: 10.1080/15548627.2023.2175569.

104. Moore BR, Page-Sharp M, Stoney JR, Ilett KF, Jago JD, Batty KT. Pharmacokinetics, pharmacodynamics, and allometric scaling of chloroquine in a murine malaria model. Antimicrobial agents and chemotherapy. 2011;55(8):3899–907. doi: 10.1128/AAC.00067-11.

