## Supplementary material for "LC3-associated endocytosis facilitates extracellular Tau aggregate internalization and degradation in microglia": SI

**Dr. Subashchandrabose Chinnathambi -** 0000-0002-5468-2129

### Supplementary figure 1

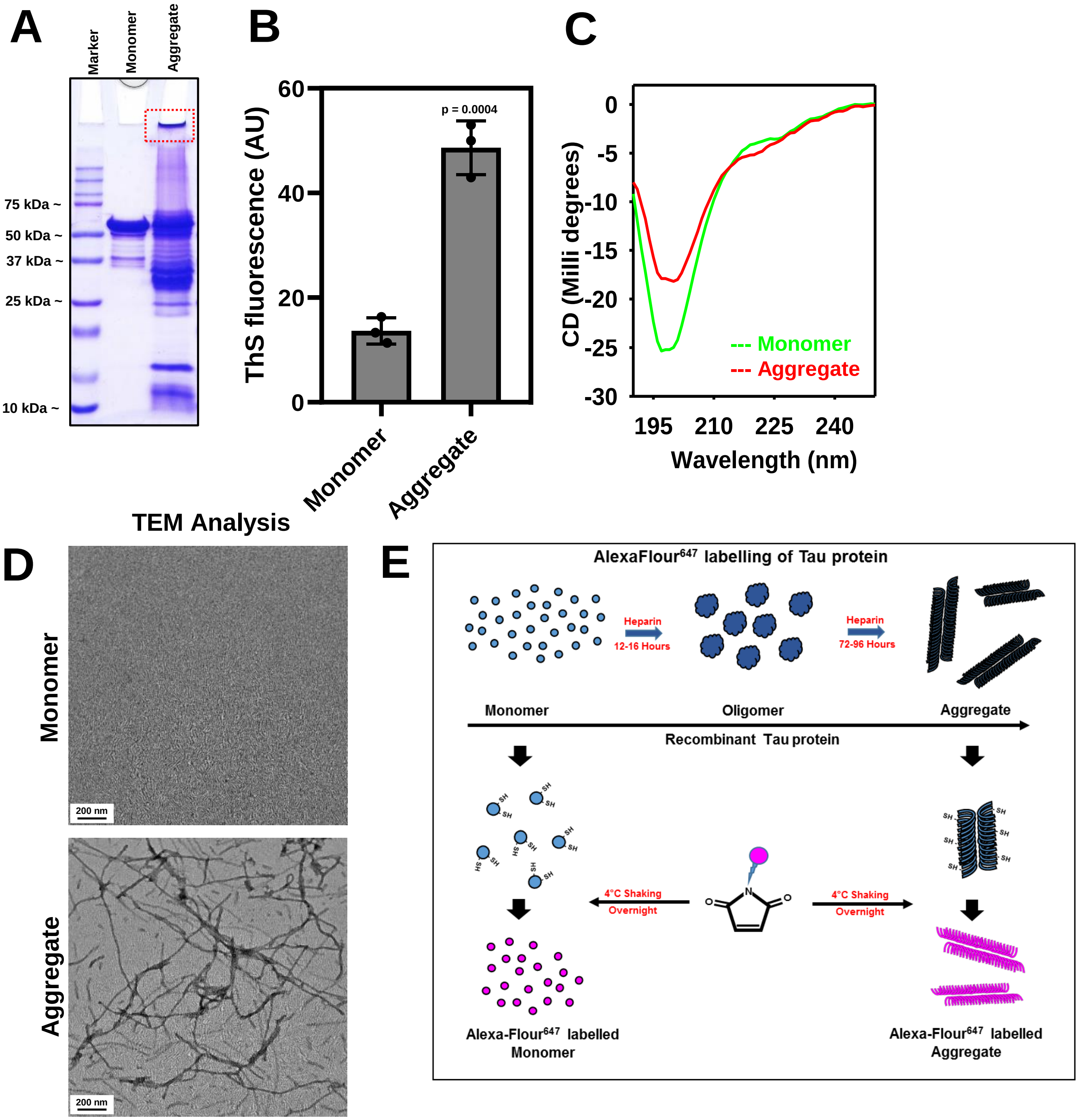


**Supplementary figure 1. Biophysical characterization and labeling of Tau.** (A) SDS-PAGE electrophoresis (10%) demonstrating monomer at ~60 kDa and SDS stable higher-order proteins in the heparin-induced aggregate sample. The band at the wells denotes filamentous aggregates that are not resolved in the electrophoresis (highlighted in red). (B) Thioflavin-S assay showing the formation of stable β-sheet structures in the aggregate group (~50 AU) as compared to monomer (~15 AU) ($n$ = 3). (C) CD spectroscopy analysis confirming the conformational change from random coil to β-sheet structure. (D) TEM analysis of Tau monomer and aggregate negatively stained with 2% uranyl acetate. (E) The schematic diagram reveals the labeling of Tau species with the C2 maleimide derivative of Alexa Fluor$^{647}$. Statistical significance was set at $P$ <0.05 by unpaired Student's $t$-test (*** $P$ <0.001, ** $P$ <0.01, * $P$ <0.05, $^{ns}$ $P$ ≥0.05). CD: Circular dichroism; PAGE: Polyacrylamide gel electrophoresis ; CD: Circular dichroism; SDS: Sodium dodecyl sulfate; TEM: Transmission electron microscope.

### Supplementary figure 2

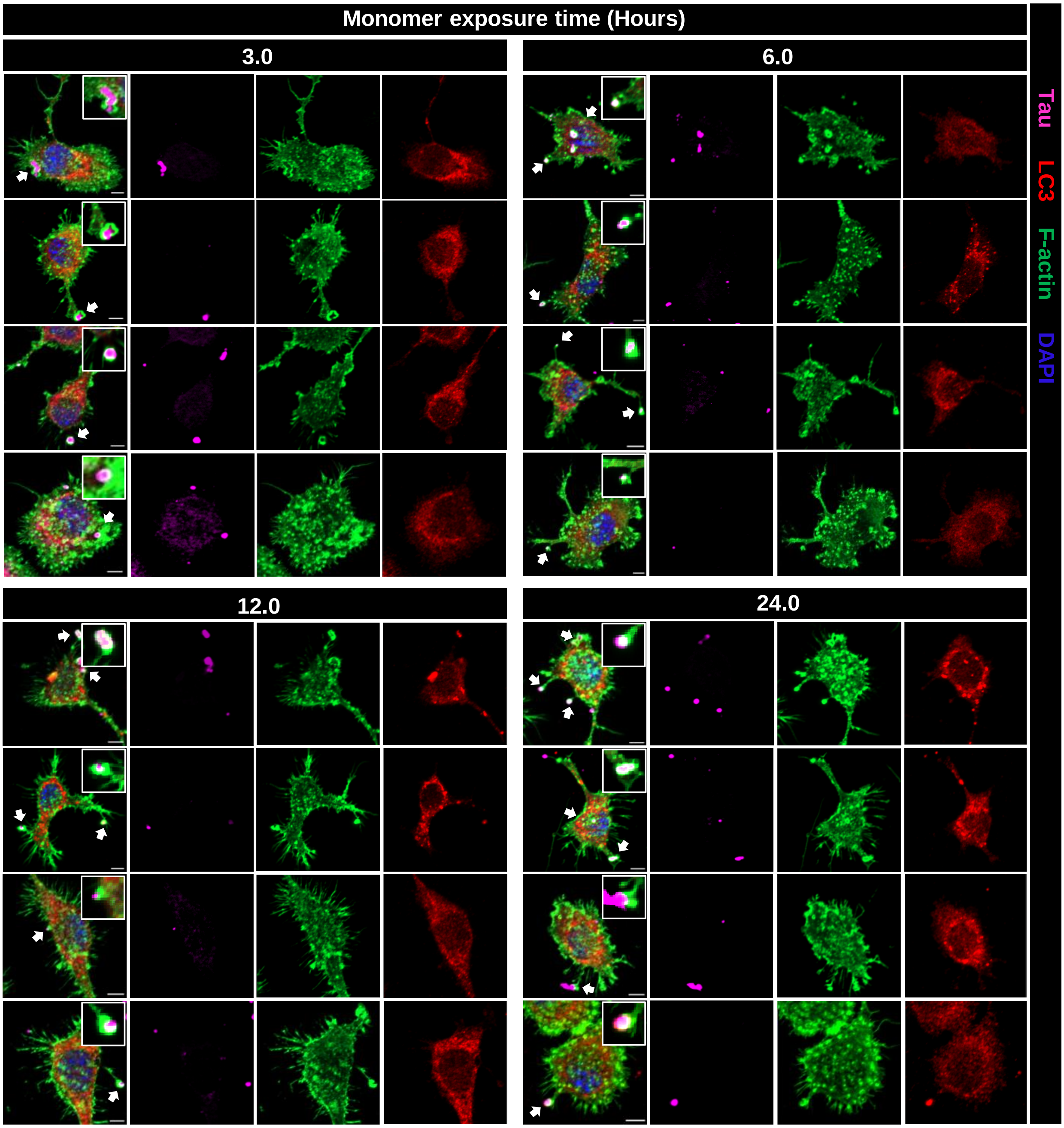


**Supplementary figure 2. Microglial endocytosis of Tau monomer.** N9 microglial cells are treated with Alexa Fluor[647]-tagged monomeric Tau (1 µM) at different time-intervals such as 3, 6, 12 and 24 hours are stained with LC3 and phalloidin (F-actin). Endosomal structures observed for monomer internalization at all time-intervals. Scale bar represents 5 µm.

### Supplementary figure 3

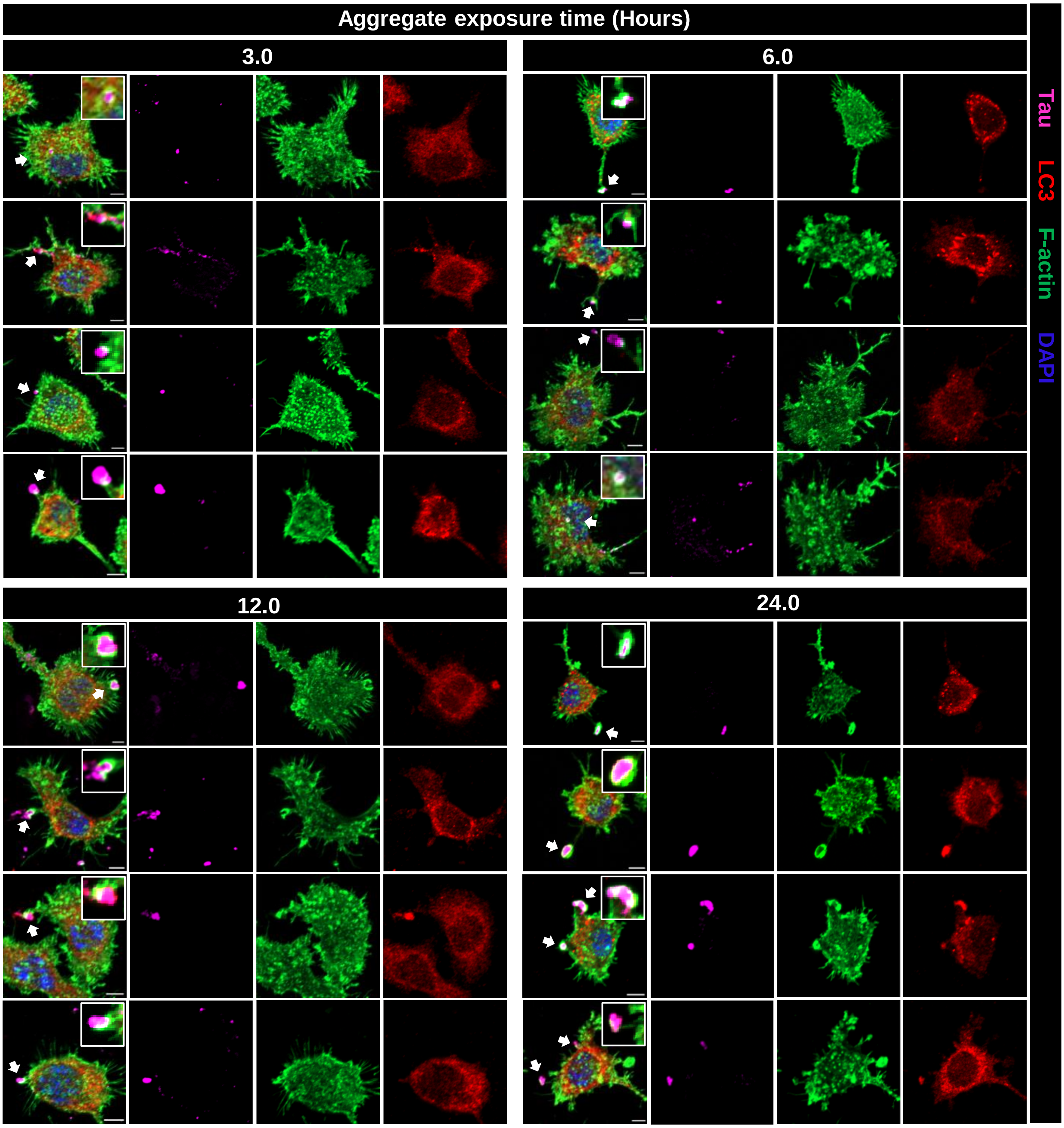


**Supplementary figure 3. Microglial endocytosis of Tau aggregate.** N9 microglial cells are treated with Alexa Fluor$^{647}$-tagged aggregated Tau (1 µM) at different time-intervals such as 3, 6, 12 and 24 hours are stained with LC3 and phalloidin (F-actin). Endosomal structures observed for aggregate internalization at all time-intervals. Scale bar represents 5 µm.

### Supplementary figure 3

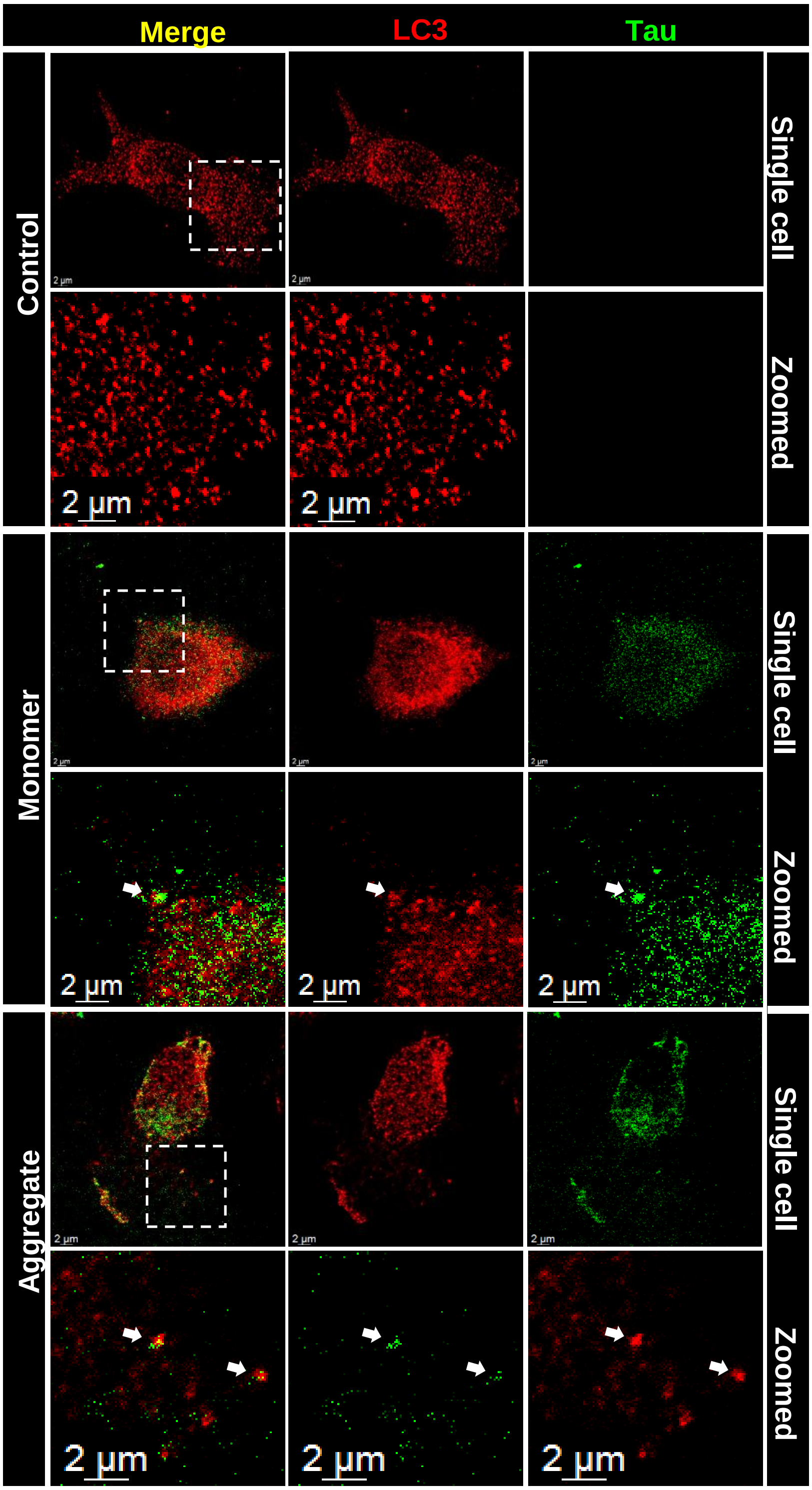


**Supplementary figure 4. LC3-associated endocytosis of extracellular Tau.** (A) Confocal microscope images of N9 microglial cells treated with Alexa Fluor[647]-tagged monomeric and aggregated Tau for 3 h stained with LC3 and phalloidin (F-actin). LC3 and Tau channels are visualized for the colocalization of LC3 with Tau (from figure 3A). The two-dimensional representation of microglial cells internalizing extracellular Tau (green) that colocalizes with LC3 (red) acquired by Leica Stellaris 5 confocal microscope (arrows indicate the endocytic structure in Tau-treated groups).
